# Decaying and expanding Erk gradients process memory of skeletal size during zebrafish fin regeneration

**DOI:** 10.1101/2025.01.23.634576

**Authors:** Ashley Rich, Ziqi Lu, Alessandro De Simone, Lucas Garcia, Jacqueline Janssen, Kazunori Ando, Jianhong Ou, Massimo Vergassola, Kenneth D. Poss, Stefano Di Talia

**Affiliations:** Department of Cell Biology, Duke University Medical Center, Durham, NC, USA; Duke Center for Quantitative Living Systems, Duke University Medical Center, Durham, NC, USA; Department of Genetics and Evolution, University of Geneva, 1211 Geneva, Switzerland; Department of Physics, École Normale Supérieure, Paris 75005, France; Duke Regeneration Center, Duke University Medical Center, Durham, NC, USA; Morgridge Institute for Research, Madison WI USA; Department of Cell and Regenerative Biology, University of Wisconsin, Madison, WI, USA; Department of Physics, University of California, San Diego, CA, USA

## Abstract

Regeneration of an amputated salamander limb or fish fin restores pre-injury size and structure, illustrating the phenomenon of positional memory. Although appreciated for centuries, the identity of position-dependent cues and how they control tissue growth are not resolved. Here, we quantify Erk signaling events in whole populations of osteoblasts during zebrafish fin regeneration. We find that osteoblast Erk activity is dependent on Fgf receptor signaling and organized into millimeter-long gradients that extend from the distal tip to the amputation site. Erk activity scales with the amount of tissue amputated, predicts the likelihood of osteoblast cycling, and predicts the size of regenerated skeletal structures. Mathematical modeling suggests gradients are established by the transient deposition of long-lived ligands that are transported by tissue growth. This concept is supported by the observed scaling of expression of the essential epidermal ligand *fgf20a* with extents of amputation. Our work provides evidence that localized, scaled expression of pro-regenerative ligands instructs long-range signaling and cycling to control skeletal size in regenerating appendages.

## Introduction

Adult teleost fish and urodele amphibians possess a remarkable capacity to regenerate entire appendages following an injury or amputation. During regeneration, a patterned, functioning structure emerges from an amputation stump months to years after its initial development. Importantly, appendage size and structure at time of loss is restored and miniaturization or overgrowth is prevented – a concept referred to as positional memory (Broussonet 1789, Morgan 1900, Poss 2010, Otsuki and Tanaka 2022). The mechanisms by which animals set the blueprint for this level of precision and read it out, often years later, during tissue regeneration has been enigmatic for centuries.

Recent work has identified potential components of positional memory. One model posits that inducer molecules are deposited in a graded fashion along the length of uninjured structures and endow positional identity to cells at homeostasis (Wolpert 1969, Adell, Cebria et al. 2010, Otsuki and Tanaka 2022). Two such proteins, Prod1 and Tig1, have been identified in salamanders: both factors are expressed in a gradient along the proximodistal axis of the salamander limb, and misexpression of either factor disrupts distal patterning (da Silva, Gates et al. 2002, Kumar, Gates et al. 2007, Oliveira, Knapp et al. 2022). In zebrafish, transcriptomic and proteomic profiling of uninjured zebrafish caudal fins identified a number of molecular candidates that are differentially expressed along the proximodistal axis (Rabinowitz, Robitaille et al. 2017). These include *aldh1a2* and *aldh1l1*, genes encoding enzymes involved in the synthesis of retinoic acid which has been known to proximalize cell fates for decades (Niazi and Saxena 1978, Maden 1982, White, Boffa et al. 1994). A non-mutually exclusive, alternative mechanism by which positional memory could be encoded is position-dependent chromatin modification. In support of this model, the level of repressive histone H3K27me3 was found to vary along the proximodistal axis of axolotl limbs, and this gradient of histone modification correlated with gene expression along the regenerating limb axis (Kawaguchi, Wang et al. 2024). However, the intracellular events that translate these molecular markers of positional memory into patterned regenerative growth remain incompletely understood.

Control of tissue growth in a regenerating adult limb or fin poses challenges distinct from those faced by developing embryonic primordia. In addition to remembering the blueprint of what has been lost, once source tissue is stimulated to regenerate, its key patterning signals must operate over large length scales (millimeters or greater) in a rapidly changing adult structure. Signaling waves have emerged as a mechanism for rapidly transporting signaling cues over long distances in regenerating tissues of zebrafish and planaria (De Simone, Evanitsky et al. 2021, Fan, Chai et al. 2023). However, it is unclear whether waves play a role in the readout of positional memory. Alternatively, scaling morphogen gradients and/or mechanical feedback, which control tissue size in the growing wing imaginal disc of *Drosophila*, could interpret positional memory during fin regeneration (Entchev, Schwabedissen et al. 2000, Tanimoto, Itoh et al. 2000, Teleman and Cohen 2000, Shraiman 2005, Kicheva, Pantazis et al. 2007, Ben-Zvi, Pyrowolakis et al. 2011, Hamaratoglu, de Lachapelle et al. 2011, Wartlick, Mumcu et al. 2011, Woodard, Li et al. 2012, Wartlick, Julicher et al. 2014). Exploring these and other mechanisms for reading out positional memory ideally requires an experimental system in which changes in biochemical signaling, cell behaviors, and mechanical forces can be followed in real time and with high sensitivity in entire cell population throughout the regeneration process. Moreover, it is important to use rigorous computational analysis to generate quantitative models that can be tested with genetic and pharmacological manipulations.

Regenerating zebrafish fins provide a compelling platform for studying positional memory. Under homeostatic conditions, the caudal fin contains 16-18 independent bony rays separated by soft interray tissue; each ray comprises multiple cell types, including epidermal cells, bone matrix-secreting osteoblasts, and mesenchymal fibroblasts. Following amputation, a wound epidermis quickly forms across the injury site and eventually secretes regeneration-promoting growth factors, including Fgf20a, that are critical for fin regeneration (Whitehead, Makino et al. 2005, Shibata, Yokota et al. 2016). Subsequently, blastemas appear at the tip of each injured fin ray and contain progenitors of skeletal structures. The growth rate of injured rays is proportional to the amount of tissue removed from the rays: Proximal amputations, which truncate rays closer to the base of the fin, remove more tissue and yield faster growth rates than distal amputations, which injure rays closer to the outgrowing fin tip. In this way regenerating rays attain their pre-injury length at approximately the same time (Lee, Grill et al. 2005, Azevedo, Sousa et al. 2012, Kujawski, Lin et al. 2014, Rabinowitz, Robitaille et al. 2017, Daane, Lanni et al. 2018, Shibata, Liu et al. 2018, Wang, Tseng et al. 2019, Uemoto, Abe et al. 2020, Ortega Granillo, Zamora et al. 2024, Surette, Donahue et al. 2024).

Here, we performed a longitudinal analysis of Erk signaling events during zebrafish fin regeneration. We chose to study this pathway for two reasons. First, it has a canonical role in regulating cell proliferation (Lavoie, Gagnon et al. 2020). Second, many previous studies have implicated Fgf and Erk signaling in directing fin regeneration (Poss, Shen et al. 2000, Lee, Grill et al. 2005, Whitehead, Makino et al. 2005, Shibata, Yokota et al. 2016). Extending these previous studies, we used tools to quantify Erk signaling sensitively and longitudinally across the entire bone-depositing, osteoblast population within regenerating fin rays. Our results show that osteoblast Erk activity persists in dynamic, long-range gradients that extend from the growing distal regenerate tip to the amputation site. Our experimental observations, combined with modeling, argue that the number of *fgf20a*-expressing epidermal cells established soon after injury instructs the formation of Erk signaling gradients in osteoblasts and that these gradients evolve in space and time due to growth-mediated transport of ligands. These Erk gradients predict the pattern of osteoblast cycling and forecast the amount of skeletal tissue that ultimately regenerates. Thus, our work elucidates how an early molecular event linked to position can be translated into a growth program that controls the size of regenerating bones.

## Results

### Osteoblast proliferation is proportional to the amount of tissue removed

Zebrafish fin rays regenerate within 3 to 4 weeks of amputation (Figure 1A). Regenerating fin osteoblasts are derived from existing osteoblasts within the ray that dedifferentiate and migrate to contribute to blastema formation at the amputation stump (Knopf, Hammond et al. 2011, Tu and Johnson 2011, Stewart, Gomez et al. 2014). A second source of joint osteoblast progenitors can also contribute to blastema formation (Ando, Shibata et al. 2017). Furthermore, the number of cells cycling within the blastema increases with the amount (severity) of amputated tissue, a cellular indicator of positional memory in this system (Lee, Grill et al. 2005). To build spatiotemporal maps of skeletal growth, we measured the expansion of osteoblast clusters located in different proximodistal regions of regenerating fins. Using transgenic zebrafish expressing histone (H2A) tagged with photoconvertible mEOS2 specifically in osteoblasts, we photoconverted and followed groups of nuclei in 4 days post-amputation (dpa) regenerates at three locations: near the amputation plane (proximal), in the middle of the regenerate (middle), and near the distal tip (distal) (Figure 1B-D). We used the edges of these clones as fiduciary marks and computed the expansion rate of the converted and non-converted regions, tiling the regenerate from distal tip to amputation plane (Figure 1C). We found that tissue expansion occurs exclusively in the distal-most 20% of regenerating rays (Figure 1E). To infer the contribution of osteoblast proliferation to ray elongation, we quantified the increase in osteoblast number in different regions of the regenerate (Figure 1F). Consistent with cell cycling being a major driver of tissue elongation, approximately 70% of the change in area occupied by photo-converted nuclei could be explained by the change in number of photo-converted nuclei (Figure 1G, solid line).

**Figure 1.**
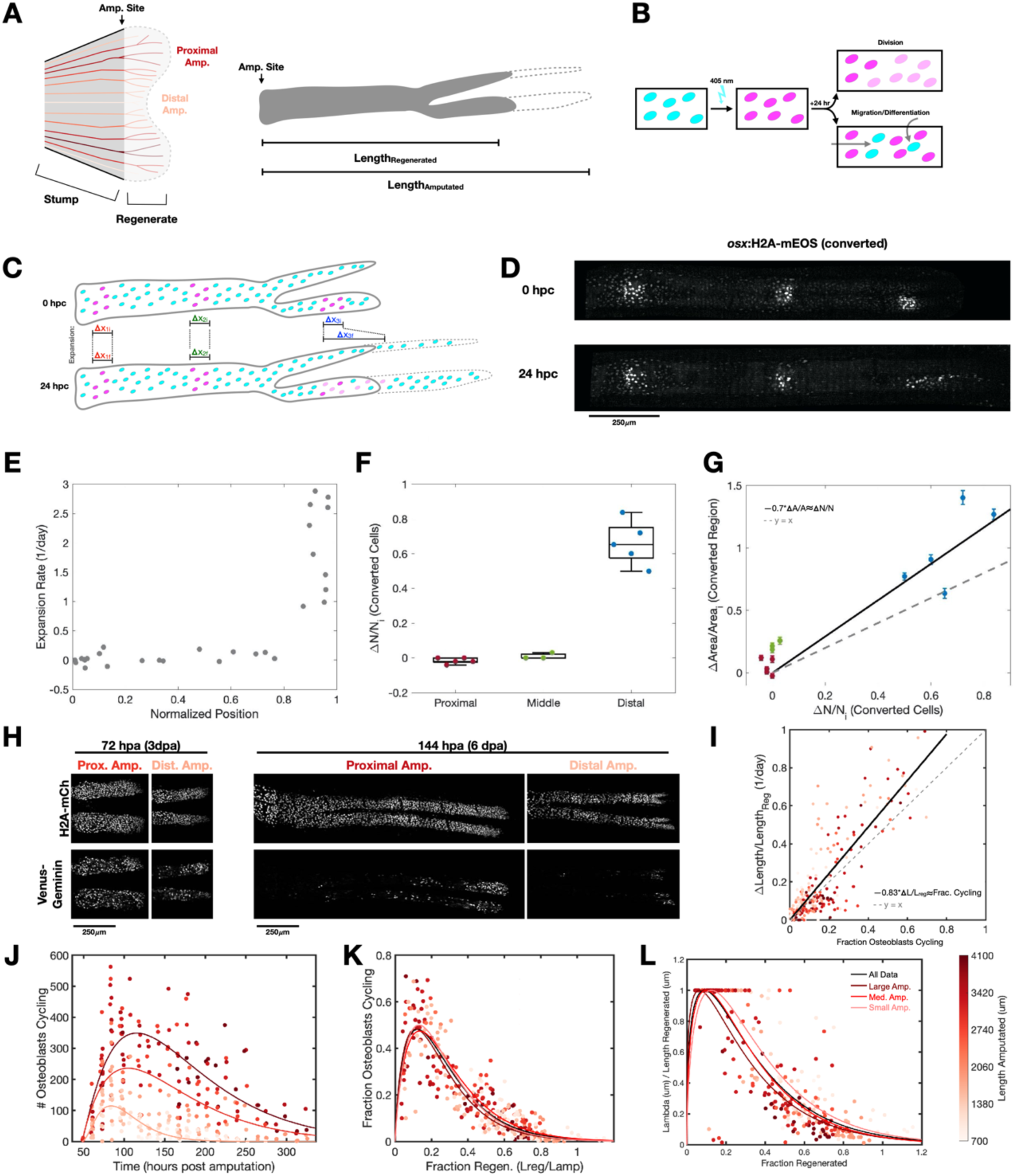
Osteoblast proliferation scales with the amount of tissue removed throughout regeneration. A) Left: Schematic of regenerating caudal fin. Rays are colored according to their length amputated. Right: Schematic of a single regenerating bony ray. B) Schematic of photoconversion experiment and outcomes expected for converted nuclei if cell division, migration, or differentiation occurs. C) Schematic of photoconversion experiment analysis. Hpc: Hours post conversion. D) Regenerating ray expressing H2A-mEOS2 specifically in osteoblasts. H2A-mEOS2 was photoconverted in three regions. The photoconverted channel is shown. E) Quantification of the expansion rate of photoconverted and neighboring unconverted regions. Normalized position is the position along the proximodistal axis divided by the length regenerated at the timepoint analyzed. Nuclei were photoconverted at 96 hpa and analyzed at 96 and 120 hpa. (Data are from 5 fish.) F) Quantification of the change in number of nuclei with photoconverted mEOS2 at regions near the amputation site (proximal), in the middle of the ray (middle), and near the distal tip (distal). (Data are from 13 clones from 5 fish.) G) Comparison of the change in area occupied by photoconverted nuclei to the change in number of photoconverted nuclei. Solid line is a linear fit of the data. Dotted line represents the identity line. Data points are colored as in Figure 1F. (Data are from 13 clones from 5 fish.) H) Representative images of regenerating rays expressing *osterix*:Venus-Geminin and *osterix*:H2A-mCherry at 3 and 6 days post amputation with larger (proximal) or smaller (distal) amputation lengths. I) Quantification of the change in ray length as a function of the fraction of osteoblasts cycling (as scored by Geminin expression). Solid line is a linear fit to the data. Dashed line is the identity line. (Data are from 60 rays from 22 fish.) J) Quantification of the number of cycling osteoblasts (as scored by Geminin expression) as a function of days post amputation. Each dot represents a single regenerating ray colored by length amputated. Curves are fits of aggregated large, medium, and small length amputated data. (Data are from 60 rays from 22 fish.) K) Quantification of fraction of cycling osteoblasts (as scored by Geminin expression) as a function of fraction regenerated (Length Regenerated / Length Amputated). Each dot represents a single regenerating ray colored by length amputated. Red curves are fits of aggregated large, medium, and small length amputated data. Black curve is fit to all data. (Data are from 60 rays from 22 fish.) L) Quantification of length of osteoblast cycling domain normalized by length of regenerate as a function of fraction regenerated. Each dot represents a single regenerating ray colored by length amputated. Red curves are fits of aggregated large, medium, and small length amputated data. Black curves are fit to all data. (Data are from 60 rays from 22 fish.)

To assay patterns of osteoblast cycling throughout the course of regeneration, we longitudinally imaged the first two weeks of regeneration in transgenic zebrafish expressing *osterix* driven RFP-tagged histone (H2A-RFP) and an *osterix* driven Venus-tagged fragment of human Geminin (Venus-Geminin) protein, which accumulates in nuclei of cycling cells (Cox, De Simone et al. 2018). We used the Venus-Geminin signal to score cycling osteoblasts, assayed by computational image segmentation of all osteoblast nuclei within regenerating rays (Amat, Lemon et al. 2014, De Simone, Evanitsky et al. 2021). During the first 3-4 days of regeneration, osteoblast cycling occurs throughout growing rays (Figure 1H, left). As regeneration progresses, Venus-Geminin expression becomes restricted to the distal tip, eventually occurring only in ∼20% of the regenerate (Figure 1H, right), consistent with the photo-conversion experiments. We found a strong correlation between the fraction of osteoblasts cycling and the rate of tissue elongation (Figure 1I). Osteoblast cycling explains ∼83% of the observed changes in osteoblast tissue length (Figure 1I), a value similar to the one estimated by our photoconversion experiments. Work in the axolotl spinal cord found that shortening of the cell cycle is important for the accelerated tissue growth seen during regeneration (Cura Costa, Otsuki et al. 2021). Conversely, we found that S/G2/M is completed in approximately 20 hours in long, short, early, and late regenerating rays (Supplemental Figure 1A). Thus, cell cycle duration does not change as a function of ray position or stage of regeneration. These observations establish that regenerative growth of osteoblast tissue is concentrated at the distal 20% of regenerating fin bones and that this growth is primarily driven by osteoblast proliferation.

We separated rays into three groups based on amputation length (large, mid-size, and small). We found that rays with larger amputated lengths contain more cycling osteoblasts throughout regeneration (Figure 1J). In contrast, the fraction of osteoblasts cycling has an invariant relationship with fraction regenerated across lengths amputated ranging from 700 *μm* or 4 *mm* (Figure 1K – black fitted curve). This suggests that the probability of osteoblast proliferation depends only on the fraction of tissue regenerated. This is due in part to the size of the proliferation domain. Indeed, the inverse relationship between the ratio of the cycling domain length to regenerate length and the fraction of tissue regenerated was independent of the amount of tissue amputated (Figure 1L). Overall, this analysis argues that the mechanism controlling osteoblast proliferation during fin regeneration involves maintaining proportionality between the fraction of cycling osteoblasts and the fraction of regeneration that remains to be completed.

### Erk activity levels encode the probability of osteoblast cycling

Extracellular signal-regulated kinase (Erk) is a canonical regulator of cell proliferation (Lavoie, Gagnon et al. 2020). Recent work from our labs identified Erk activity as a regulator of timed bursts of osteoblast hypertrophy in regenerating zebrafish scales (De Simone, Evanitsky et al. 2021). To test if Erk activity patterns osteoblast cycling during fin ray regeneration, we treated animals with the MEK inhibitor PD03 for 24 hours at 4 dpa. During this treatment, rays of DMSO-treated fish added significantly more tissue (131 *μm* on average) than rays of PD03-treated fish (7 *μm* on average) (Figure 2A, B). Furthermore, the number of cycling cells did not measurably change in rays of DMSO-treated fish, whereas the number of cycling cells decreased by 78% in rays of PD03-treated fish (Figure 2C, D). Thus, Erk inhibition blocked skeletal growth and osteoblast cycling.

**Figure 2.**
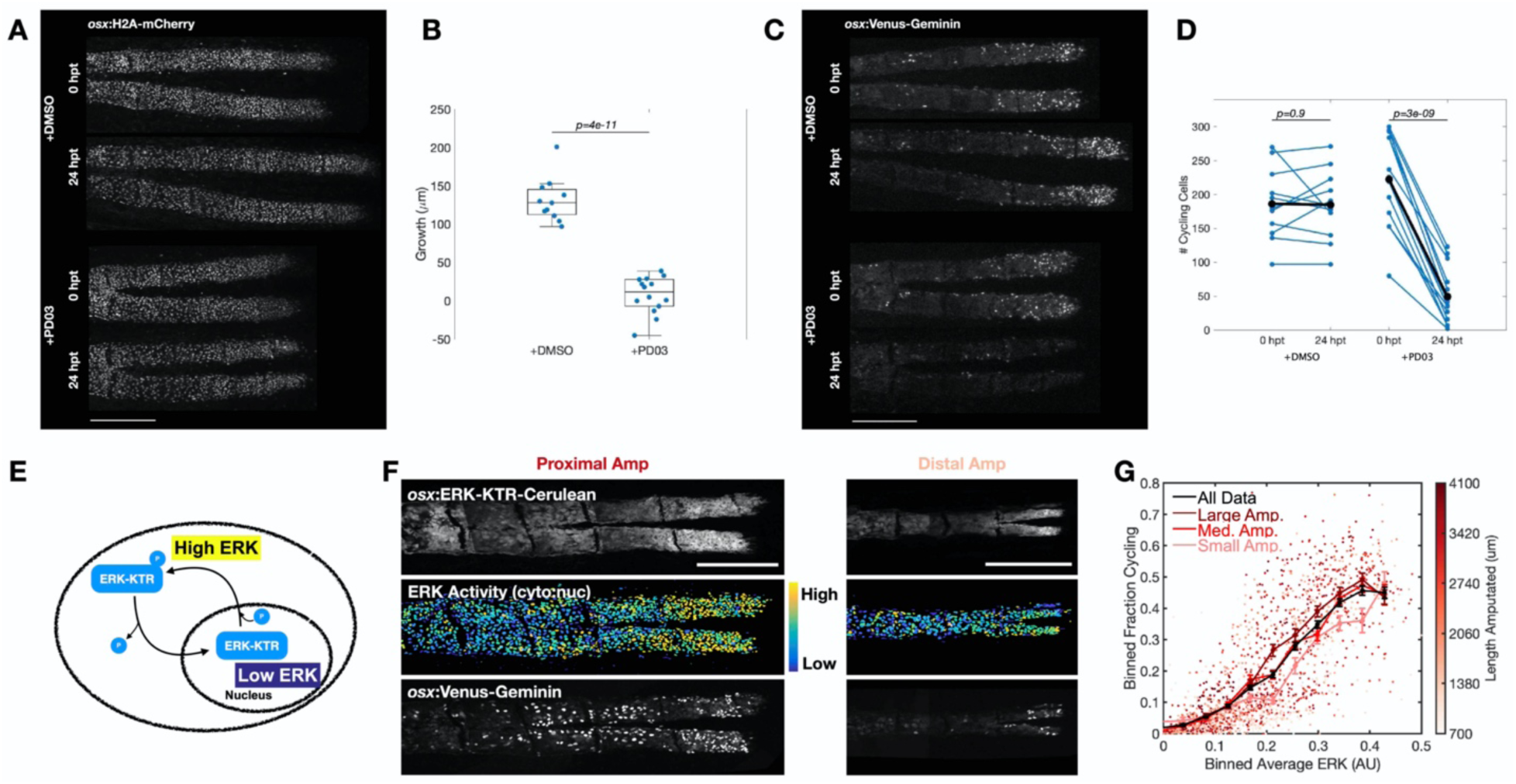
Erk activity encodes the probability of cell cycling. A) Representative image of a ray from a fish expressing *osx*:H2A-mCherry treated with control DMSO (top) or MEK inhibitor (PD03, bottom). Scale bar is 250 *μm*. B) Quantification of change in length of osteoblast tissue following 24 hours of treatment with a MEK inhibitor (PD03). (Data are from 9/12 untreated fish/rays and 10/14 treated fish/rays. Two sample t-test was used to test for significant difference.) C) Representative image of a ray from a fish expressing *osx*:Venus-Geminin treated with control DMSO (top) or MEK inhibitor (PD03, bottom). Scale bar is 250 *μm*. D) Quantification of the number of osteoblasts cycling (as scored by Venus-Geminin expression) before and after MEK inhibition. Black dots and lines in indicate average values. (Data are from 8/11 untreated fish/rays and 10/14 treated fish/rays. Paired t-test was used to test for significant difference.) E) Schematic describing the biosensor used to measure Erk activity in regenerating osteoblasts. F) Representative images of regenerating rays with large (proximal) and small (distal) lengths amputated expressing the Erk biosensor (top), Venus-Geminin (bottom), and Histone2A-mCherry (not shown) in osteoblasts. Heatmap (middle) has Erk activity values for individual osteoblasts plotted onto the nuclei of those osteoblasts, as segmented on the Histone2A-mCherry channel. Color indicates single-cell Erk activity level. Scale bars are 250 *μm* G) Quantification of the relationship between fraction of osteoblasts cycling and average Erk activity. Each dot represents a spatial bin occupying 1/10^th^ of the regenerate’s proximodistal axis, time averaged over a 24 hour window. (See methods and Supplemental Figure 1F.) Color indicates length amputated. Black line represents binned average of all data. Red lines represent binned average of aggregated large, medium, and small length amputated data. (Data are from 1869 regions/20 fish/55 rays.)

To map Erk dynamics during fin regeneration, we longitudinally imaged a fluorescently-tagged Erk kinase translocation sensor that reports Erk activity as the ratio of cytoplasmic to nuclear sensor fluorescence (Figure 2E, F) (Regot, Hughey et al. 2014, De Simone, Evanitsky et al. 2021). Previously, traveling waves of Erk activity that pattern bursts of osteoblast hypertrophy were identified in regenerating zebrafish scales (De Simone, Evanitsky et al. 2021). To determine whether waves of Erk activity also exist in regenerating fin bones, we imaged the same rays every 8 hours for 2-3 days (Supplemental Figure 1B). We detected noisy temporal oscillations of Erk activity with a typical period of ∼24 hours (Supplemental Figure 1C, D). However, these oscillations were correlated only over short distances of about 200 *μm* and did not give rise to long range patterns, as would be expected for a traveling wave (Supplemental Figure 1E). Thus, unlike regenerating zebrafish scales, osteoblasts in regenerating fins do not display detectable traveling waves of Erk activity.

To quantify the relationship between osteoblast cycling and Erk activity, we imaged regeneration in fish co-expressing the Erk biosensor and Venus-Geminin. To account for the oscillatory nature of Erk activity and potential temporal delays between Erk activity and Geminin expression, we compared spatiotemporally averaged Erk activity values and osteoblast cycling (see methods for details, Supplemental Figure 1F). We found that regions of regenerating rays with higher average Erk activity possess higher probabilities of osteoblast cycling (Figure 2G). For example, regions of regenerating rays with average Erk activities of 0.4 AU versus 0.08 AU have osteoblast cycling probabilities of 46% and 5%, respectively. This quantitative relationship between Erk activity and osteoblast cycling is similar regardless of how much tissue is amputated (Figure 2G). To test this relationship, we titrated Erk activity with increasing amounts of the MEK inhibitor (0 – 10 µM PD03). We found that intermediate concentrations of PD03 decrease Erk activity in regenerating osteoblasts and yield proportionally reduced amounts of osteoblast cycling (Supplemental Figure 1G).

### Erk activity patterns explain growth dynamics observed during fin bone regeneration

To determine if tissue regeneration, in addition to osteoblast cycling, can be predicted by Erk activity levels, we quantified Erk activity in all osteoblasts of individual regenerating rays, and for each ray we calculated average Erk activity levels across the whole regenerate. By plotting these against fraction regenerated or time post amputation (Figure 3, Supplemental Figure 2), we found that average osteoblast Erk activity levels decrease (from ∼0.4 AU to 0 AU) in fin rays over the course of regeneration (Figure 3C, Supplemental Figure 2B, C, E). This decrease is almost linear with respect to fraction regenerated (Figure 3C – black line), and the linear relationship is the same even when we analyze separately groups of rays with small, mid-size, or large lengths amputated (Figure 3C – colored lines). This observation, that average Erk activity has an invariant relationship with fraction regenerated across at least a 6-fold range of amputation lengths, together with the finding that Erk controls osteoblast cycling, suggests that average Erk activity is a crucial regulator of the dynamic patterning of cell cycle decisions in regenerating osteoblasts.

**Figure 3.**
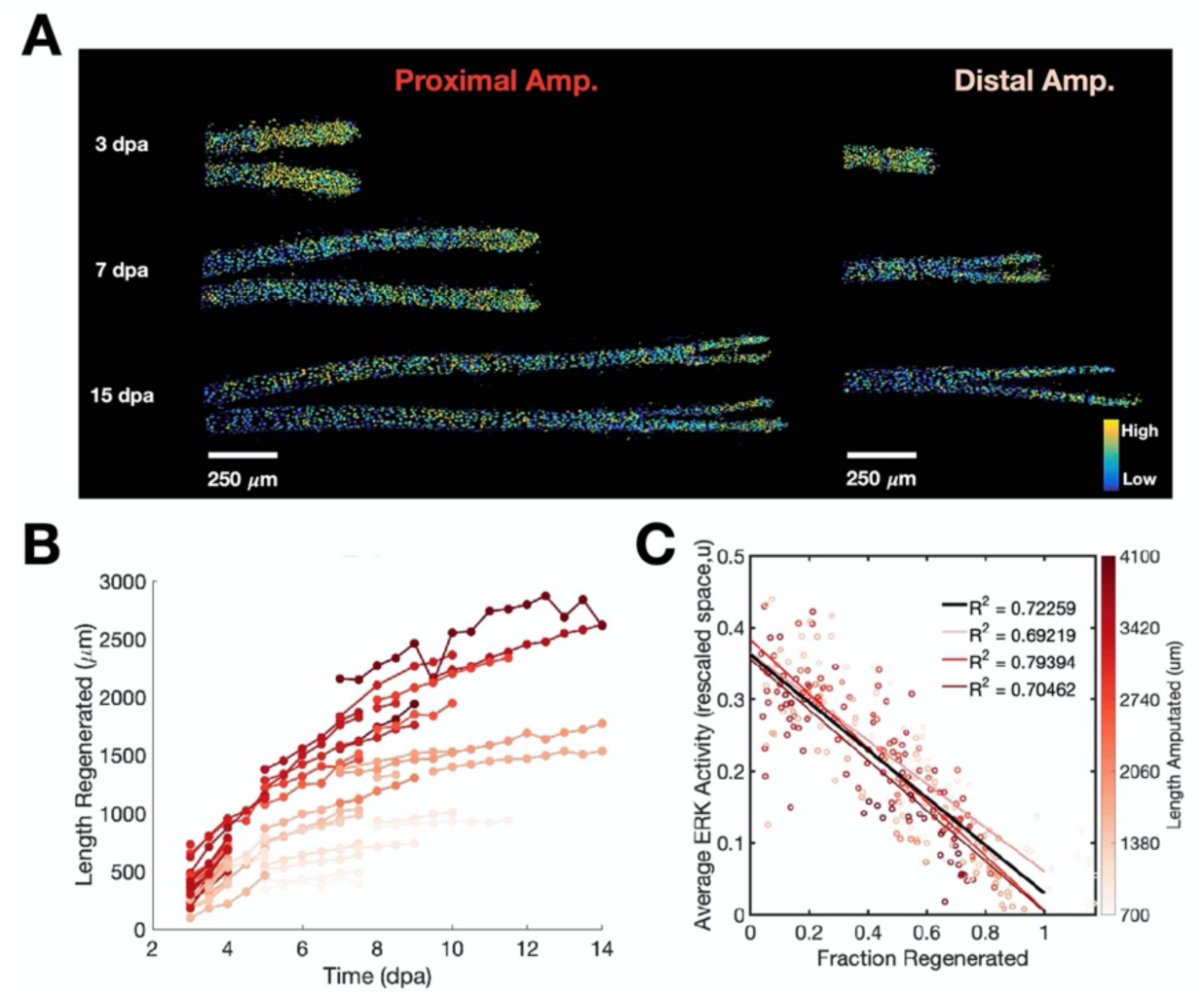
Whole-regenerate averaged Erk activity explains regenerative outgrowth. A) Representative heatmaps of Erk activity in regenerating rays throughout the first 2 weeks of regeneration from proximal (left) and distal (right) amputations. B) Quantification of regenerate length over time for all analyzed rays. Lines connect data points from the same ray. Coloring indicates length amputated. Data are from 60 rays from 22 fish. C) Whole ray average Erk activity versus fraction regeneration (Length Regenerated / Length Amputated). Each dot represents a single ray at a single time point, colored according to length amputated. Red lines indicate fits to data sub-grouped by amputation length. Black line indicates fit to all data. Data are from 60 rays from 22 fish.

To map proximodistal patterns of osteoblast Erk activity, we analyzed Erk levels along the length of the regenerate (Figure 4A-B). Over time, Erk activity decreases at all proximodistal positions (Figure 4B). To account for this decrease and still compare Erk patterns at different times during regeneration, we normalized local activity measurements within each ray by the average Erk activity over the whole regenerating ray (Erk activity / Whole Regenerate Average Erk Activity) and plotted these measurements as a function of normalized position (Regenerate Position / Regenerate Length) (Supplemental Figure 3C). These normalizations enable the extraction of spatial profiles of Erk activity regardless of regenerate length or Erk activity amplitude. To filter out the noise in spatial profiles arising at each time point from the oscillatory nature of Erk activity (Supplemental Figure 1B-D), we computed the spatial Erk activity pattern of individual rays by time-averaging data over 24-hour windows. This time window approximates both the typical Erk oscillation period and cell cycle duration of proliferating cells (Supplemental Figure 1A, D). We computed such averages across all imaged times here and throughout the manuscript, unless otherwise noted. We found that the normalized spatial Erk activity maps of all 60 individual rays imaged formed gradients stretching from distal tip to amputation site (Figure 4C-D, Supplemental Figure 3D-E). The similarity of these gradients is striking as they were obtained from rays with lengths amputated ranging from 700 *μm* to 4 *mm* and across post-amputation times ranging from 3 to 14 days (Figure 4C-D, Supplemental Figure 3D-E).

**Figure 4.**
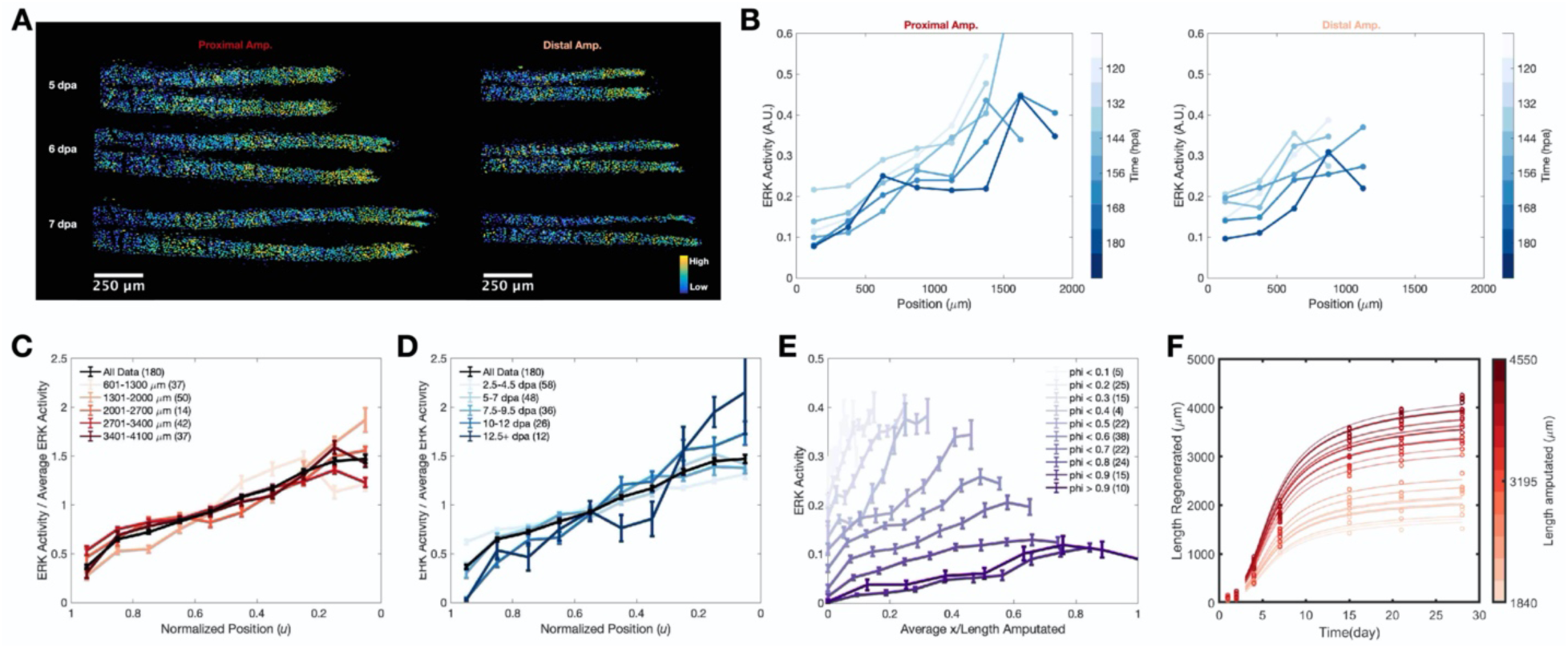
Erk activity is patterned in scaling gradients that span the entire regenerate and encode proliferation across space. A) Representative images of Erk activity heatmaps for regenerating rays with proximal (left) or distal (right) amputations, imaged every 12 hours from 5 to 7 days post amputation. B) Quantification of Erk activity along the proximodistal axis the regenerating rays shown in Figure 4A. Amputation site is plotted at 0 on x-axis. Each dot represents the average Erk activity of cells within a spatial bin. Lines connect points from the same ray at a given time. Coloring indicates time post amputation. C) Quantification of normalized average Erk activity (Binned Average Erk Activity / Whole Ray Average Erk Activity) versus normalized position along the proximodistal axis (position / Length Regenerated). Each red line represents the average spatial Erk Activity profile of a group of rays with the indicate lengths amputated. Black line represents the average spatial Erk activity profile for all rays analyzed. Data are from 52 rays from 20 fish. D) Quantification shown in Figure 4C with data grouped by time post amputation instead of length amputated. Data are from 52 rays from 20 fish. E) Representative spatiotemporal profile of Erk Activity throughout the course of osteoblast regeneration. Each line represents the average spatial Erk Activity profile for rays of the indicated fraction regenerated. All rays were first divided into 10 equal spatial bins and binned average Erk activity was calculated for each spatial bin. These binned values were then averaged for each spatial position across all rays in a given phi group. Each curve was plotted as a function of normalized position in a region spanning from the amputation plane to the median phi value of a given phi group. F) Comparison of the fin ray growth dynamics predicted by our quantitative model (lines) versus an independent experimental characterization of ray growth dynamics (open circles). Colors indicate length amputated.

Plotting the gradients at different times as a function of position normalized by length amputated (after grouping rays with similar fractions regenerated), provides an intuitive understanding of the dynamics of the Erk gradients (Figure 4E). The gradients initiate with high maximum Erk activity (∼0.4 AU) and are very steep. Over time they both extend and flatten out so that their length scale matches the regenerate length and the ratio of Erk activity at the tip and the amputation plane is unchanged (∼4-fold). The amplitude of the gradient (defined as the value at the tip) decreases as regeneration proceeds, from a high amplitude (0.4 AU) to a negligible amplitude (<0.1AU), at which time tissue growth stops (Figure 4E). In addition, we performed a series of analyses of the properties of Erk activity values, concluding that our data support a model in which oscillations of similar amplitude across space are overlaid on long-range gradients (Supplemental Figure 4A-D).

Collectively, our data reveal an invariant relationship between average Erk activity and fraction of tissue regenerated (Figure 3C) and an invariant spatial gradient of normalized Erk activity (Figure 4C-D). Average Erk values across the whole regenerate decrease, reflecting the flattening of gradients over time. Additionally, the Erk gradients stretch, effectively scaling with regenerate length over time. Thus, for any normalized position along the proximodistal axis, Erk activity decreases similarly in all regenerates, regardless of length amputated. Moreover, regardless of the amount of tissue amputated, when a fin ray reaches a certain fraction regenerated (for example, 0.4 or 0.6), the whole-regenerate averaged Erk activity (∼0.24 or ∼0.16 AU) is proportional to the remaining fraction of length amputated that must be replaced (that is 0.6 or 0.4). We therefore propose that the gradients of osteoblast Erk activity act as rulers that measure past growth and determine future growth.

Our quantitative observations on Erk activity allow us to decouple the gradient into a length and an amplitude component. The length of the gradient scales with the length of the regenerate, while its shape is invariant across all fish, rays, and times. The amplitude of the gradient decays over time as the fraction of tissue regenerated increases, essentially encoding the stage of regeneration and the amount of tissue that remains to be regenerated. This can be written mathematically as:

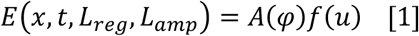

where *x* indicates the position within the regenerate, *t* indicates time post amputation, *L_reg_* indicates the length of the regenerate, *L_amp_* indicates the length amputated, *φ* = *L_reg_*/*L_amp_*, and *u* = *x*/*L_reg_*. Here, *A*(*φ*) describes the behavior of whole-regenerate averaged Erk activity shown (Figure 3C), and *f*(*u*) describes the scaling gradients for all rays and fish at all time points (Figure 4C-D, Supplemental Figure 3D-E). We note that *f*(*u*) is derived from time-averaged data which average out the oscillatory Erk dynamics. This ansatz, Eq. [1], is a major simplification of Erk dynamics as it brings a four variable dependency down to two variables. Moreover, the relationship shown in Eq. [1] is factorized. That is, it depends on the product of two functions, each dependent on one variable only, further separating the control of Erk activity temporally and spatially.

To test the hypothesis that Erk activity gradients encode osteoblast growth dynamics, we used this mathematical description of Erk activity and the relationship between Erk activity and cycling (Figure 2E) to predict tissue growth. Assuming that growth is driven by proliferation, we obtain:

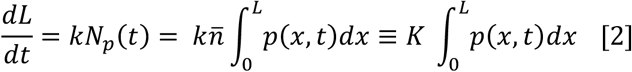

where *L* is the length of the regenerate, *N_p_* is the number of proliferating cells, *ñ* is the average density of osteoblasts along the ray length (which we found to be almost constant in space), and *p*(*x*, *t*) the probability of proliferation at position *x* and time *t*. We note that *p*(*x*, *t*) is defined so that its spatial average 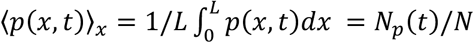 coincides with the fraction of proliferating cells. In normalized spatial coordinates, *u* = *x*/*L*,this equation becomes:

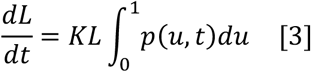

Approximating the dependency of proliferation on Erk activity (*E*) with a power-law, i.e. *p*(*u*, *t*) = *βE^α^*(*u*, *t*), and using Eq. [1] to describe Erk, we obtain:

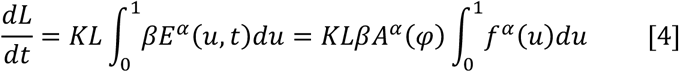

By dividing the left- and right-hand sides of this equation by *L_amp_* and by defining 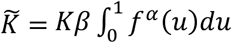, we obtain:

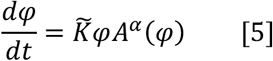

This implies that the growth of the ray depends only on the amplitude of the gradient. The gradient shape is integrated out, as 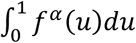 is the same for all rays at each time point. Since all the parameters in Eqs. [4-5] can be experimentally estimated, we predicted tissue growth in a fit-independent manner. We found good agreement between the growth dynamics predicted by the quantitative model and the dynamics experimentally observed in separate, independent experiments measuring ray growth dynamics (Figure 4F). Thus, our quantitative model of how Erk activity patterns osteoblast cycling in space and time recapitulates growth dynamics observed during fin bone regeneration.

### Erk dynamics can explain regenerative growth dynamics of *longfin* mutants

To further test the importance of Erk activity in regulating size control during regeneration, we asked whether the relationship between Erk activity and tissue growth persists in fins of *longfin* (*lof*) fish (Iovine and Johnson 2000), genetic mutants that overexpress a voltage-gated potassium channel, *kcnh2a*, and overgrow rays in homeostatic and regenerative contexts (Figure 5A, B and Supplemental Figure 5A-C) (Geraudie, Monnot et al. 1995, Stewart, Le Bleu et al. 2021). We visualized and measured Erk activity and Venus-Geminin expression in individual regenerating *lof* osteoblasts throughout the first three weeks of regeneration (Figure 5C). Using these measurements and the same computational approach as in Figure 2E, we determined that the relationship between Erk activity and the probability of osteoblast proliferation is similar in *lof* and wildtype fish (Figure 5D). We note that rays of *lof* fish were imaged every 48 hours, versus every 12 hours for wildtype. Spatial profiles of Erk activity in *lof* fish were, therefore, not time averaged. Furthermore, as in wildtype fish, Erk activity is patterned in long-range gradients throughout regeneration in *lof* osteoblast tissue, regardless of length amputated or time post amputation (Figure 5E & Supplemental Figure 5D). Similar to wildtype rays (Figure 3C), Erk activity decreases linearly throughout regeneration in *lof* rays (Figure 5F), but there is not a universal relationship between whole-regenerate averaged Erk activity and fraction regenerated in *lof* rays of different lengths (Figure 5F). This is consistent with the fact that size memory is compromised in *lof* fish, with ventral lateral rays undergrowing and medial rays overgrowing relative to their amputated length (Figure 5B and Supplemental Figure 5C) (Geraudie, Monnot et al. 1995). Instead, we found a universal, nearly linear relationship between whole-regenerate averaged Erk activity and actual fraction regenerated (Length Regenerated/Final Length Regenerated) in *lof* fish (Figure 5G). Thus, Erk activity correlates with the actual length regenerated in *lof* bony rays, consistent with Erk activity instructing growth. This relationship between Erk activity and osteoblast cycling spans a 5-fold range of lengths amputated (Figure 5G), indicating scaling akin to that observed in wildtype animals (Figure 3C). Thus, Erk activity remains predictive of osteoblast cycling and growth dynamics in a mutant where size memory is compromised.

**Figure 5.**
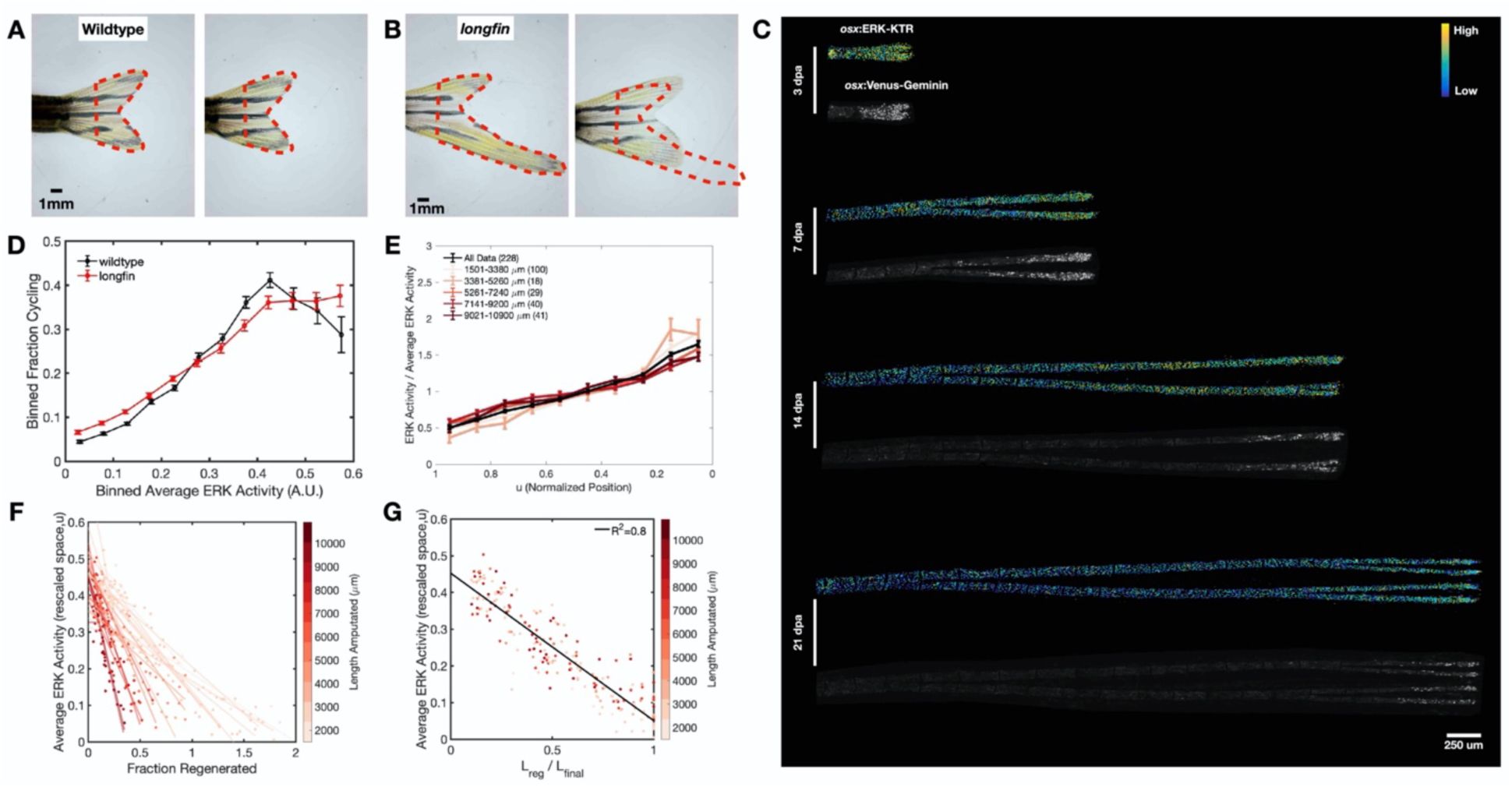
Erk dynamics can explain regenerative growth dynamics of *longfin* mutants. A,B) Representative images of regenerating wildtype (A) and *lof* (B) fins before (left) and after (right) amputation. Red dotted line indicates fin shape before amputation. C) Representative Erk activity heatmaps and images of Venus-Geminin expression in a single regenerating *lof* ray over the first three weeks of regeneration. D) Comparison of the relationship between Binned Erk Activity and Fraction Osteoblasts Cycling (as scored by Venus-Geminin expression) in wildtype (black) and *lof* (red) fish. LOF data are from 29 rays from 7 fish. WT data are from 60 rays from 20 fish. E) Spatial Erk activity profiles of *lof* rays, as in Figure 4C. Individual rays were grouped into 5 groups based on their length amputated. The average spatial Erk Activity profile of each group is shown in red. The average spatial Erk Activity profile of all rays is shown in black. Data are from 36 rays from 9 fish. F) Quantification of whole ray average Erk activity versus fraction regenerated (length regenerated / length amputated). Each dot represents a single ray at a single time point, colored by length amputated. Lines indicate fits to data from individual rays throughout regeneration. Data are from 36 rays from 9 fish. G) Quantification of whole ray average Erk activity (as in Figure 5F) versus actual fraction regenerated (length regenerated / final regenerated length measured). Each dot represents a single ray at a single time point, colored by length amputated. Black line indicates fit to all data. Data are from 36 rays from 9 fish.

### Growth-induced advection can explain long-range Erk activity gradients

To test if Fgf signaling, a canonical activator of Erk activity, instructs Erk gradient formation, we treated fish transiently with BGJ398, a pan-FgfR inhibitor (Figure 6A). As expected, based on many studies implicating Fgf signaling in fin regeneration (Poss, Shen et al. 2000, Lee, Grill et al. 2005, Whitehead, Makino et al. 2005, Shibata, Yokota et al. 2016), we found Erk activity decreased on average by 62% in the rays of BGJ-treated fish but only by 7% in rays of DMSO- treated control fish (Figure 6B). The number of cycling osteoblasts decreased by 95% in rays of BGJ-treated fish but increased by 14% in rays of DMSO-treated control fish (Figure 6C). Consequently, osteoblast tissue of treated but not control fish stopped elongating following pan-FgfR inhibition (Fig 6A, D). Thus, as expected, Fgf signaling is required for Erk activity and Erk-dependent cycling in osteoblasts.

**Figure 6.**
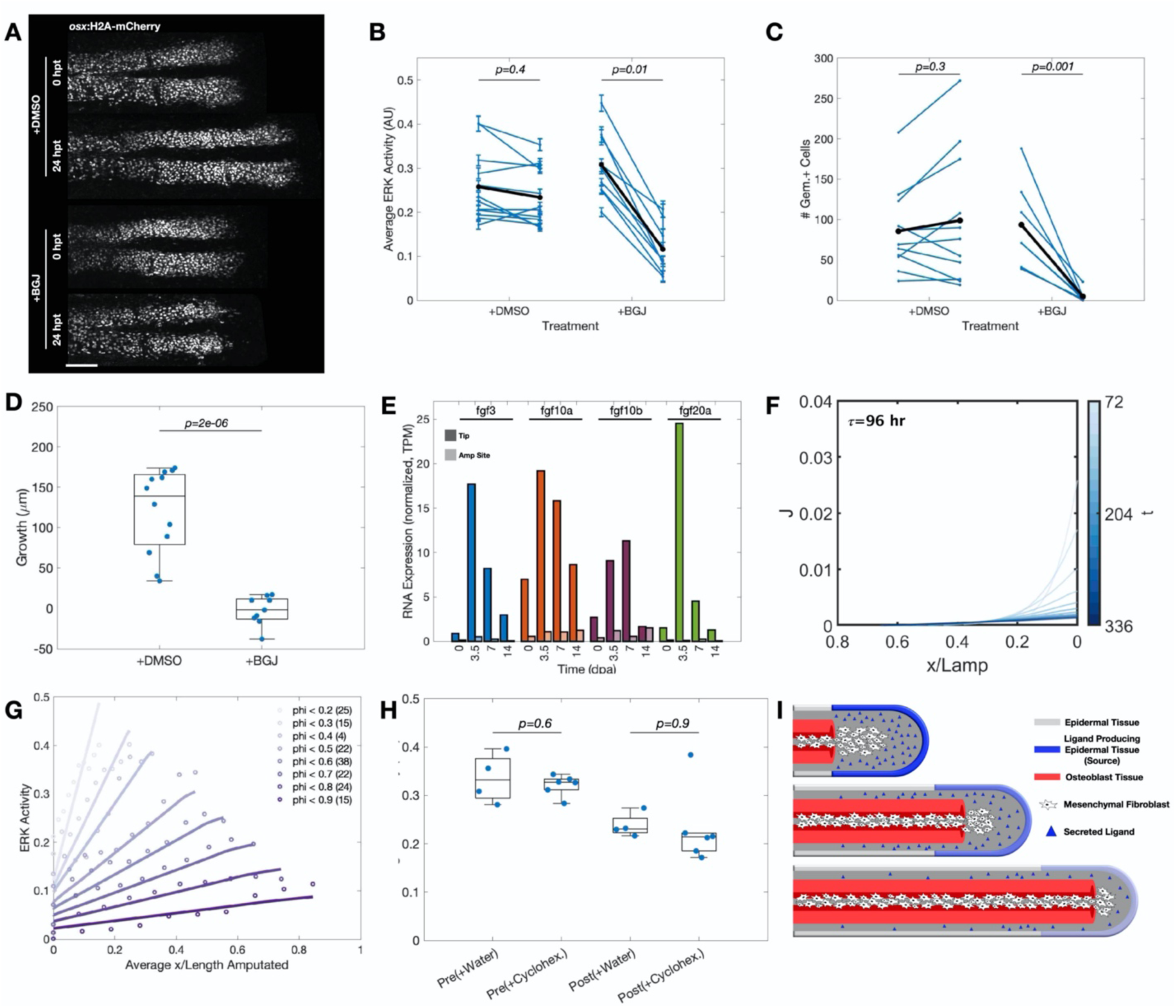
Long-range Erk activity gradients require Fgf signaling and likely form by tissue expansion-induced advection. A) Representative images of regenerating rays of fish expressing *osterix*:H2A-mCherry treated with DMSO (top) or a pan-FGF receptor inhibitor (BGJ, bottom) for 24 hours. B, C, D) Quantification of Erk activity (B), number of cells expressing Venus-Geminin (C), and tissue growth (D) in DMSO and pan-Fgf receptor inhibited (BGJ-treated) fish. Black dots and lines in (B & C) indicate average values. Data are from: B – 11 untreated rays from 7 untreated fish & 8 BGJ-treated rays from 6 BGJ-treated fish. C&D – 12 untreated rays from 8 untreated fish & 9 BGJ-treated rays from 7 BGJ-treated fish. E) Bulk RNA sequencing of FGF ligand expression at the tip (dark bars) and amputation site (light bars) at 0, 3.5, 7, and 14 days post amputation. See methods for details. F) Predicted ligand-producing source dynamics derived from theoretical model using 96 hour ligand lifetime. Color indicates time. G) Comparison of computationally predicted and experimentally observed spatiotemporal profile of Erk Activity throughout the course of osteoblast regeneration. Experimental data (circles) are repeated from figure 4E. Simulated data are shown in lines. H) Quantification of Erk activity in untreated and translation-inhibited (cycloheximide treated) fish before (left) and after 12 hour treatment (right). Data are from 4 untreated rays from 4 untreated fish and 6 cycloheximide treated rays from 6 cycloheximide treated fish. I) Schematic depicting advection-dilution model of Erk activity gradient formation. Early in regeneration, a transient ligand-producing source is established in the distal epidermis (blue). This domain produces and secretes long-lived ligand (blue triangles), which activates Erk in osteoblasts (red). Tissue growth advects and dilutes the ligand, yielding Erk activity gradients that span the regenerate from distal tip to amputation plane.

We next investigated how Fgf signaling directs Erk gradient formation and maintenance by developing a mathematical model of the system. Four processes likely contribute to the observed gradients: ligand production, ligand diffusion, transport of cells and ligands by tissue growth, and ligand processing (degradation). Approximating fin rays as a 1D system leads to the following mathematical model:

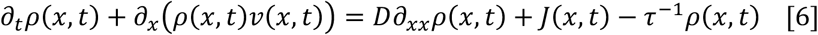

where *ρ* is the density of ligands, *v*(*x*, *t*) is the velocity field describing cellular flows, *D* is the diffusion constant of the ligand(s), *J*(*x*, *t*) is the source, or rate of ligand production, and *τ* is the lifetime of the ligand. To simulate this model, we need to evaluate these parameters/functions. Based on our previous experiments on cell proliferation and its dependence on Erk (Eqs. 2-4), we can express the velocity field as:

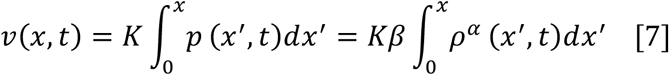

where we have used the previously assumed power-law dependence of proliferation on Erk activity.

Next, we estimated the spatiotemporal properties of the ligand source by performing bulk RNA-sequencing of tissue isolated from the distal tip and amputation plane. We detected expression of genes encoding several Fgf ligands known to be expressed during fin regeneration at the distal tip, including *fgf20a, fgf3,* and *fgf10a* (Shibata, Yokota et al. 2016, Cudak, López-Delgado et al. 2023). These genes showed a 16-, 20-, and 3-fold increase, respectively, in expression at the distal tip compared to 0 dpa (Figure 6E). This expression generally peaked at 3.5 days post amputation and returned to baseline by 14 days post amputation. We did not detect substantial expression of any Fgf ligand near the amputation plane at any time (Figure 6E). These results indicate that Fgf ligands are produced transiently and specifically near the distal regenerate tip.

To determine potential mechanisms that can produce the observed experimental gradients, we first considered the role of diffusion. However, several arguments detailed in the Methods argue against a major role for diffusion given the characteristic lengths and timescales of the observed gradients. Second, we considered the possibility that the gradients form through a physical mechanism involving both transport of ligands by tissue growth (a process known as advection) and ligand processing/degradation. This corresponds to solving Eq. [6] with *D* = 0. To determine if this model can reproduce the experimentally observed Erk gradients, one can use standard numerical methods to compute *ρ*(*x*, *t*) for a given source *J*(*x*, *t*) and lifetime of the ligand *τ*. However, these parameters are difficult to evaluate quantitatively, so we took advantage of our description of Erk dynamics and tissue flows (Eqs. [1] and [7]) to determine which spatiotemporal dynamics of the source and of the ligand lifetime reproduce the experimentally observed gradients (see Methods for details). We found that a transient and localized source of ligands (Figure 6F) can generate the dynamic, scaling gradients of Erk activity observed experimentally (Figure 6G), as long as the lifetime of the ligand is ≥ 4 days. In other words, for the model to hold the ligand would need to be relatively stable and not rapidly degraded. To test the plausibility of a long lasting Fgf activating ligand, we treated fish with cycloheximide to block protein synthesis and quantified the decay rate of Erk activity. If ligand processing is very fast, we expect Erk activity to rapidly disappear upon cycloheximide treatment. In contrast, we found that Erk activity changed similarly in fish treated with cycloheximide for 12 hours versus control fish (Figure 6H). Longer cycloheximide treatments are toxic, but this result suggests the lifetime of ligand processing is at least a few days. We note that several processes may impact this lifetime, including ligand-receptor interactions, ligand internalization and degradation, and receptor expression.

This model suggests a simple mechanism for the scaling of the gradients and the control of growth. The only length scale in the problem is the spatial extension of the source (Figure 6F). Thus, if the source size is proportional to the amount of tissue amputated, Erk gradients will be expanded proportionally, proliferation will be scaled, and tissue growth will be proportional to the amount of tissue removed, directing accurate skeletal replacement (Supplemental Figure 6). This must be contrasted with scaling in a diffusion-degradation model, where the length scale of the gradient is given by 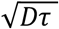. This would require that within the same animal the product of *D* and *τ* changes by 2 orders of magnitude in a fine-tuned manner, since we observe gradients scaling over at least one order of magnitude in length: short gradients (∼200 *μm*) exists in medial rays early in regeneration while long gradients (∼2 − 4 *mm*) exist late in regeneration in lateral rays.

Figure 6I diagrams intuitively how an advection-dilution model could distribute a long-lived ligand along the length of the regenerate and yield the long-range Erk signaling gradients we observe in regenerating osteoblast tissue. A transient ligand-producing source at the tip of the regenerate leaves behind a trail of ligands as the tissue grows. If these ligands are sufficiently long-lived, the gradients can extend across the entire tissue and will be scaled by the growth dynamics and size of the source to match the length of the regenerate. Thus, we argue that the advection-dilution model presented here provides a robust physical mechanism for controlling the formation and expansion of signaling gradients in large growing tissues.

### Fgf20a expressing epidermal tissue scales with length amputated, suggesting a mechanism of positional memory

An advection-dilution model requires that the ligand-producing source scales with length amputated during fin osteoblast regeneration. We therefore sought experimental evidence for this critical condition. While the identity of the Fgf ligand that activates signaling during osteoblast regeneration is not experimentally confirmed using conditional genetics, Fgf20a is essential for blastema formation and can induce expression of other Fgf ligands (Whitehead, Makino et al. 2005, Shibata, Yokota et al. 2016). Furthermore, *fgf20a*-expressing epidermal cells that directly overlie regenerating osteoblasts are in the appropriate place at the right time to influence Erk activity in regenerating osteoblasts (Figure 7A) (Shibata, Yokota et al. 2016). Consequently, Fgf20a was employed here as a representative ligand to assess the validity of our theoretical model.

**Figure 7.**
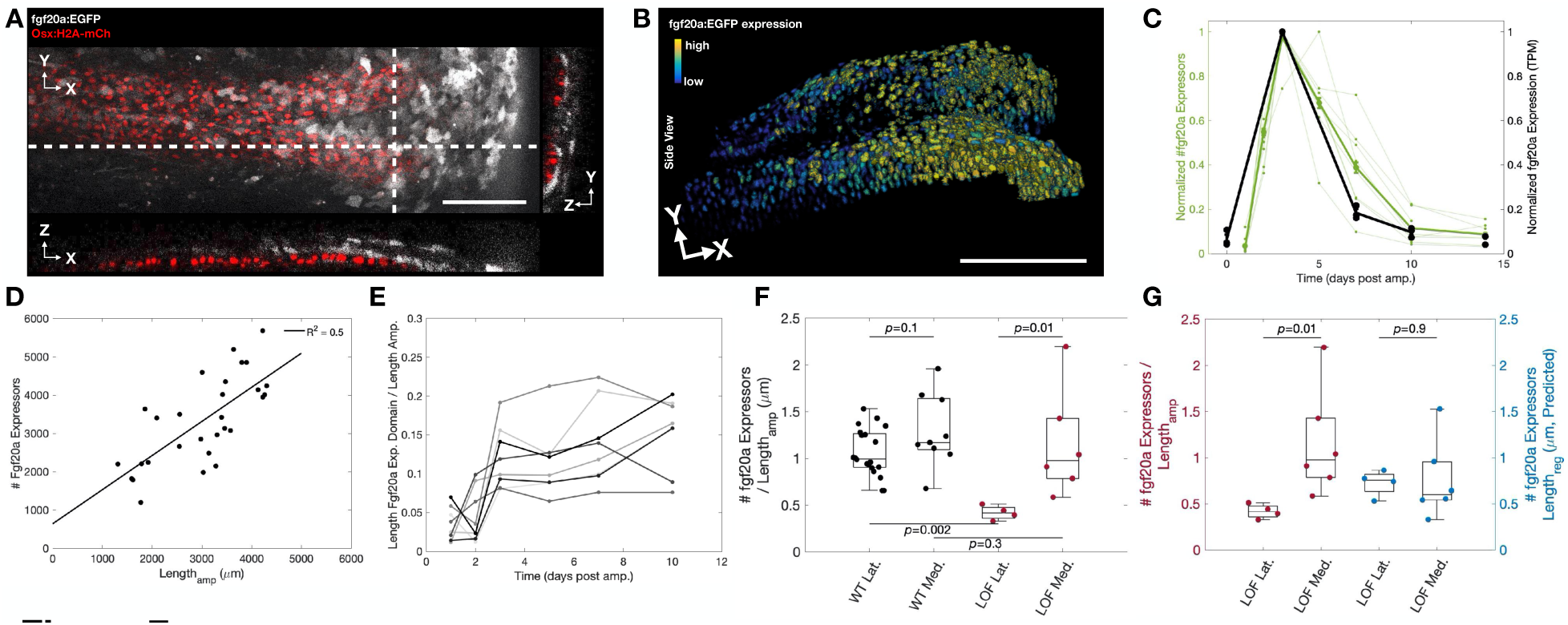
The number of *fgf20a* expressing epidermal cells scales with length amputated suggesting a mechanism of positional memory. A) Representative image of regenerating ray expressing GFP driven by *fgf20a* regulatory sequences (*fgf20a*:EGFP, gray scale) and *osterix*:H2A-mCherry (red). Dotted lines indicate regions shown as YZ and XZ projections. Scale bar is 100 *μm*. B) Quantification of *fgf20a:*EGFP expression projected in 3D as heatmap onto epidermal nuclei (segmented on *krt19*:H2A-mCherry expression, not shown). Scale bar is 100 *μm*. C) Quantification of *fgf20a:*EGFP expression versus time post amputation. The number of *fgf20a:*EGFP expressing cells was scored at 1, 2, 3, 5, 7, 10, and 14 days post amputation (green data). Plotted data are normalized to the maximum number of *fgf20a:*EGFP expressing cells detected at any time point. Thin green lines are data from individual regenerating rays. Bold green line is average of all regenerating rays. Bulk RNA expression of Fgf20a was assayed at 0, 3, 7, 10, and 14 days post amputation (black data). Plotted data are normalized to values at 3 dpa. See methods for RNAseq details. Data for *fgf20a:*EGFP expression are from 30 rays from 12 fish. D) Quantification of the number of epidermal cells expressing *fgf20a:*EGFP at 72 hpa and the length of tissue amputated. Each dot represents a single regenerating ray. Line represents fit to all data. Data are from 26 rays from 12 fish. E) Quantification of the length of the *fgf20a:*EGFP expression domain normalized by length amputated versus time post amputation. Lines connect data from single regenerating fish and are colored in gray scale to distinguish individual rays. Data are from 8 rays from 4 fish. F) Quantification of the ratio between the number of epidermal cells expressing *fgf20a:*EGFP divided by the ray length amputated for lateral and medial wildtype (black) and *lof* (red) rays. Wildtype data are from 26 rays from 12 fish. lof data are from 10 rays from 4 fish. G) Quantification of the ratio between the number of epidermal cell expressing *fgf20a:*EGFP divided by the predicted regenerate length of *lof* lateral and medial rays (right, blue data). Ratio between the number of epidermal cells expressing *fgf20a:*EGFP and the length amputated of *lof* lateral and medial rays is repeated from Figure 7F for reference (left, red). Data are from 10 rays from 4 fish.

To test whether the size of the Fgf20a production domain scales with the length of tissue amputated, we live-imaged a transgenic line co-expressing *krt19:H2A-RFP* and the enhancer trap *fgf20a:*EGFP. These lines label epidermal nuclei and drive GFP expression from *fgf20a* regulatory elements, respectively. This enables us to segment individual epidermal nuclei and measure a proxy for amounts of *fgf20a* expressed by individual epidermal cells (Figure 7B). We found that the number of *fgf20a*:EGFP expressing cells peaks at 3 dpa, decreases rapidly between 3- and 7 dpa, and approaches pre-injury expression levels by 10 dpa (Figure 7C). *fgf20a*:EGFP expression dynamics observed by live-imaging are similar to the dynamics we reported in our bulk RNA sequencing experiment (Figure 7C, black line). The slight delay (∼1 day) in the rate of *fgf20a*:EGFP expression decay between live-imaging experiments and bulk mRNA-sequencing likely reflects GFP persistence. We focused on the peak *fgf20a* expression time point at 3 dpa (Figure 7C) and found a strong correlation between the number of cells expressing *fgf20a*:EGFP and the length of tissue amputated (Figure 7D). The length of the *fgf20a*:EGFP expression domain also correlates with the length of tissue amputated (Supplemental Fig 7A). These observations agree with a crucial prediction of our mathematical model: The number of source cells and the length of the source region scale with the amount of tissue removed. We further interrogated how the spatial pattern of *fgf20a*:EGFP expression changes throughout regeneration. Once established over the first 3 dpa, the size of this source domain remains constant, occupying only the distal 10-20% of the length amputated (Figure 7E). Altogether, these data indicate that the ligand producing source is transient, decaying over a few days. Furthermore, the source localizes to the distal-most region of the growing regenerate and scales in size with the amount of tissue amputated, consistent with the predictions made by our modeling.

If the growth dynamics of regenerating fin rays are set by scaling the ligand production domain, source scaling should be aberrant in *lof* mutant fish, which have compromised size memory (Figure 5B, Supplemental Figure 5C, & Supplemental Figure 7B). To test this hypothesis, we measured the correlation between the amount of tissue removed and the number of *fgf20a:EGFP* expressing cells in *lof* fish. On average, one *fgf20a*:EGFP expressing epidermal cell is established for every micron of tissue amputated in wildtype lateral and medial rays (Figure 7F). A similar relationship between number of *fgf20a*:EGFP expressing cells and length amputated is observed in the medial rays of *lof* fish (Figure 7F). In contrast, on average, only 0.4 *fgf20a*:EGFP expressing epidermal cells are produced for every micron of tissue removed from *lof* ventral lateral rays (Figure 7F), which could explain why *lof* lateral rays have a smaller fraction regenerated than medial *lof* rays (Supplemental Figure 7B). Consistently, the number of *fgf20a*:EGFP expressing epidermal cells per micron of final tissue regenerated is not statistically different between lateral and medial *lof* rays (Figure 7G, blue). Thus, while the establishment of the ligand production domain is compromised in *lof* ventral lateral rays (Figure 7F), the number of *fgf20a*:EGFP expressors specified is still predictive of regenerative outgrowth (Figure 7G). This suggests that the compromised size memory in these *lof* rays is due to poorly scaled ligand production.

We next compared regenerative outgrowth to the number of the *fgf20a*:EGFP producing cells in wildtype and *lof* fish. Wildtype lateral and medial rays generate ∼1 *μm* of tissue per *fgf20a*:EGFP expressing epidermal cell in the source domain; *lof* lateral and medial rays produce ∼1.5 *μm* of regenerate per *fgf20a*:EGFP expressing epidermal cell (Supplemental Figure 7C). Thus, while the number of *fgf20a*:EGFP expressing cells correlates with how much tissue is regenerated regardless of amputation length and ray position in both genetic backgrounds, *lof* rays exhibit more regenerative outgrowth than wildtype rays for ligand-producing source domains of a given size. This ∼1.5-fold overgrowth per *fgf20a*:EGFP expressing cell of *lof* mutants versus wildtype is not due to a longer-lived source in *lof* fish, as *fgf20a* expression decays similarly (or possibly slightly faster) in *lof* fish (Supplemental Figure 7D). In contrast, we found that Erk activity decays slower in *lof* fish (Supplemental Figure 7E), suggesting the overgrowth of *lof* mutants could be due to slower ligand decay in *lof* fish. We estimated ligand lifetimes of ∼3.8 and ∼6 days for WT and *lof* mutants, respectively, from the rate of decay of Erk activity near the amputation plane (Supplemental Figure 7E, inset). We then performed simulations of our model using different ligand processing lifetimes (Supplemental Figure 7F). When we input the ligand lifetime values estimated from wildtype (black) and *lof* (red) data into our simulations, we predicted that *lof* mutants would overgrow by about a factor 1.5x relative to WT for a given source size (Supplemental Figure 7F), which is consistent with our experimental observations.

Taken together, our data reveal that gradients of Erk activity in osteoblasts form downstream of Fgf ligand expression and influence the probability of cell cycling. Mathematical modeling suggested that these gradients arise because Erk-induced osteoblast cycling drives tissue growth that advects (transports) Erk-activating ligand away from the production domain. Quantitative imaging showed that the *fgf20a:*EGFP expression domain is proportional to the amount of tissue amputated. Thus, larger amputations yield larger domains of *fgf20a* production, which yield more total Erk activity, osteoblast cycling, and tissue growth. Ultimately, this ensures both scaling gradients and proper size control of regenerated tissue. Thus, we infer that Erk activity levels, set by ligand-expressing epidermal cells, represent a basis for positional memory in the osteoblasts of regenerating zebrafish fins.

## Discussion

Here, we propose a mechanism for how cells within a regenerating appendage encode and decode positional memory, dynamically monitoring in real time the extent of structural replacement. Using *in toto* live-imaging in combination with computational and theoretical approaches, we generated maps of Erk activity at multiple timescales and across the entire osteoblast space in regenerating fins. Our experiments reveal the presence of signaling gradients and how they contribute to processing positional memory during regeneration. Our theoretical work argues that advection, that is the bulk movement of cells and ligands due to tissue growth, is likely the physical process that drives the formation of these Erk gradients. Finally, our work suggests that formation of Fgf ligand expression domains in proportion to the amount of tissue amputated could be a key step in endowing positional memory to regenerating fin rays.

Fgf signaling has been long recognized as a key regulator of zebrafish fin regeneration (Poss, Shen et al. 2000, Lee, Grill et al. 2005, Whitehead, Makino et al. 2005, Shibata, Yokota et al. 2016). However, how Fgf signaling controls cellular decisions in space and time throughout regeneration remains incompletely understood. Waves of signaling activity have emerged as a mechanism for controlling regeneration across large spatial scales. (De Simone, Evanitsky et al. 2021, Fan, Chai et al. 2023). Alternatively, morphogen and/or metabolic gradients have long been proposed to pattern body and appendage axes during regeneration (Morgan 1905, Child 1941, Otsuki and Tanaka 2022). To determine whether signaling waves, gradients, or an alternative mechanism patterns zebrafish fin regeneration, we focused on quantifying Erk activity, a major downstream effector of the Fgf pathway. This approach revealed spatiotemporal patterns of this signaling pathway and dissected how these patterns instruct cell proliferation and tissue growth. Specifically, we discovered that Erk activity gradients, formed downstream of Fgf activity, act as rulers during fin bone regeneration. These Erk rulers inform on the amount of regenerative outgrowth that has occurred and still needs to occur. Notably, the length of each gradient is scaled (proportional) to the amount of tissue amputated, allowing for the same pro-growth signals to precisely control outgrowth regardless of length amputated. Furthermore, because these gradients develop over time and always span the length of the regenerate, they provide a mechanism of long-range coordination across the entire tissue. This eliminates the need for additional active, long-range communication, which would be difficult to control accurately across the millimeters of osteoblasts that separate the distal tip and amputation plane. Thus, we propose that the gradients we have described here provide a mechanism to accurately translate positional memory into regenerative growth. Since the mechanism we uncovered only requires a growing tissue and a long-lived ligand, we speculate that similar expanding and decaying gradients might control regeneration in other organisms and/or biological contexts.

While our discovery of Erk gradients in osteoblasts provides an understanding of how positional memory is decoded during regeneration, the mechanism that encodes this memory, upstream of gradient formation, remains incompletely understood. Our results suggest that the early establishment of an Fgf20a production domain that is scaled to the length amputated is a key step. This hypothesis is consistent with recent work that reported that fin size memory can be reset only in the earliest stages of fin regeneration (Wang, Tseng et al. 2019). If correct, this mechanism elegantly endows positional memory in a way that is robust to perturbations at later stages of regeneration and does not require long-range cell-cell communication throughout regeneration. In the future, it will be critical to identify the molecular and cellular mechanisms underlying the formation and scaling of the source domain and manipulate its properties. The earliest step in caudal fin regeneration is wound healing. In zebrafish, mechanical waves propagate through the caudal fin epidermis within hours of amputation. The distance over which these waves propagate correlates with the amount of tissue removed (De Leon, Wen et al. 2023). These mechanical waves depend on the presence of reactive oxygen species (ROS), whose levels correlate with the amount of tissue removed (De Leon, Wen et al. 2023). However, these waves have only been characterized for the first few hours of regeneration, whereas formation of the *fgf20a* expressing domain in the wound epidermis takes 2-3 days; thus, mechanisms likely exist to link these two processes and establish the population of cells expressing *fgf20a*. After wound healing, fibroblasts and osteoblasts dedifferentiate below the amputation plane and migrate across the amputation plane to form the regenerative blastema (Knopf, Hammond et al. 2011, Sousa, Afonso et al. 2011, Stewart and Stankunas 2012). Secreted immune factors regulate injury-induced osteoblast migration during this early phase of regeneration (Sehring, Mohammadi et al. 2022). It is possible that a combination of mechanical and morphogenetic signals direct cellular migration in proportion to the amount of tissue removed to sculpt the ligand-producing source domain.

Continued progress on understanding how positional memory is encoded and decoded during fin regeneration will benefit from the quantitative approach we employed here. For example, our *in toto* imaging combined with computational image analysis revealed noisy temporal oscillations of Erk activity on short time scales (8-12 hours) that averaged out to robust gradients of Erk activity at longer times scales (≥24 hours). These signaling patterns would go undetected if assessed by snapshots in end-stage imaging experiments. Additionally, our quantitative, longitudinal data allowed us to derive an analytical framework to describe Erk activity across the entire regenerative process and to derive a mathematical model that closely approximates the signaling patterns and growth dynamics of all rays independently of the amount of tissue removed. This model revealed that tissue growth itself likely plays a major role in distributing Erk-activating Fgf ligands upstream of Erk gradient formation. Notably, tissue growth was proposed to underlie long-range Fgf ligand gradient formation during vertebrate embryonic tailbud elongation (Dubrulle and Pourquie 2004).

Surprisingly, the feasibility of our model for fin regeneration requires that the activating Fgf ligand(s) in this system have a relatively long lifetime of few days. While our model makes no assumptions and provides no insight on the mechanism underlying long ligand lifetime in this system, several possibilities exist. For example, internalized ligand might persist inside cells and later be recycled (Romanova-Michaelides, Hadjivasiliou et al. 2022). Alternatively, extracellular molecules, such as heparin, could stabilize secreted Fgf ligand. This has been documented for Fgf2 in cultured neurospheres, where Fgf2 persists for days in the presence of heparin (Caldwell, Garcion et al. 2004). It will be important to test this theoretical model of Erk gradient formation and probe the lifetime of Fgf ligands in this system, ideally by directly visualizing Erk-activating Fgf ligands *in vivo* during fin regeneration.

Overall, this work, together with a growing body of work (Rompolas, Deschene et al. 2012, Tornini, Puliafito et al. 2016, Lukonin, Serra et al. 2020, Cura Costa, Otsuki et al. 2021, De Simone, Evanitsky et al. 2021, Xin, Gallini et al. 2024), demonstrates how *in toto* live imaging approaches, rigorous computational image analysis, and mathematical modeling can uncover new biological concepts and patterns in regeneration involving different molecular factors, tissues, and species.

## Methods

### Zebrafish husbandry

Wildtype or transgenic zebrafish of the outbred Ekkwill strain ranging in age from 3 to 18 months old were used in all experiments. Approximately equal sex ratios were used. Fish were housed at 4-7 fish per liter in Pentair Aquatic Habitat Recirculating Multi-Rack Systems and fed 2-3 times daily. Water temperature was maintained between 26 and 28.5°C, and fish were kept on a 14:10 hour light:dark cycle. For caudal fin amputations, fish were anesthetized in 0.75% v/v 2-phenoxyethanol (Sigma-Aldrich) in fish water and fins were amputated with a razor blade. All experiments using zebrafish were approved by the Institutional Animal Care and Use Committee at Duke University. Transgenic lines used in this study include Tg(*osx:H2A-mCherry*)^pd310^ (Cox, De Simone et al. 2018), Tg(*osx:ErkKTR-mCerulean*)^pd2001^ (De Simone, Evanitsky et al. 2021), Tg(*osterix:Venus-hGeminin*)^pd271^ (Cox, De Simone et al. 2018), Tg(*osx:H2A-mEos2*)^pd2002^(De Simone, Evanitsky et al. 2021), *fgf20a*:EGFP (Nagayoshi, Hayashi et al. 2008, Shibata, Yokota et al. 2016), *Tg(krtt1c19e:H2A-mCherry)^pd309^*(Han, Chen et al. 2019), and *longfin (lof)^d2^* mutant (Iovine and Johnson 2000).

### Live Imaging

For *in vivo* confocal imaging, adult zebrafish were anaesthetized in 0.01% tricane (VWR, MSPP-MS222) in system water and transferred to a 1% agarose bed in an acrylic glass plate, as previously described (Cox, De Simone et al. 2018). The caudal fin was placed on a glass slide and secured with a thin layer of cooling 1% agarose while the head remained partially submerged in diluted tricane. Additional cooling 1% agarose was applied to the trunk and the areas of the platform dorsal and ventral to the trunk to secure the fish. Fish were then immersed in diluted tricaine. Gill movements were monitored for the duration of imaging. When gill movements slowed, fresh system water was applied directly to the mouth with a peristaltic pump (Cole Parmer; no. EW-73160-32; silicon tubing: Tygo, 0.7 mm inner diameter and 2.4 mm outer diameter, B-44-4X) until regular gill movement was restored. Fish were returned to system water immediately following imaging. During longitudinal imaging, fish were returned to the aquatic system between time points.

Confocal imaging was performed on a Leica SP8 confocal microscope using a HC FLUROTAR L 25X/0.95 W VISIR water immersion lens (Leica 15506374) at 0.75X zoom. For each regenerate, multiple overlapping z-stacks were acquired, tiling the entire osteoblast tissue. Images were acquired at 1,024 x 1,024 resolution (0.606 μm pixel size) with a z-step of 0.606 µm. Fluorescent proteins were imaged with laser of the indicated wavelengths: Histone-mCherry (561 nm), ErkKTR-Cerulean (458 nm), Venus-hGeminin (514 nm), unconverted H2A-mEos2 (488 nm), converted H2A-mEos2 (561 nm), *fgf20a:*EGFP (488 nm). Variable laser powers were used according to transgenic reporter expression. To photoconvert mEos2, a region containing unconverted nuclei was repeatedly scanned with 405nm light at 2% laser power until nuclear conversion was observed.

Low-magnification, whole fin images were acquired on a Zeiss AxioZoom V16 using ZEN Pro 2012 (blue edition) software. Fish were anaesthetized in 0.75% v/v 2-phenoxyethanol in system water and subsequently transferred to a Petri dish and partially submerged in 0.75% 2-phenoxyethanol for imaging. Fish were returned to the aquatic system following imaging.

### Pharmacological treatments

For pharmacological treatments, fish were immersed in the pharmacological compound diluted to working concentration in fish water for the duration of the experiment. The pan-FgfR inhibitor, BGJ398 (Selleck Chem 2183), and MEK inhibitor, PD0325901 (Selleck Chem S1036), were dissolved in DMSO and used at 10 *μm* unless otherwise indicated. Control fish were treated with equivalent amounts of DMSO vehicle. The translation inhibitor, cycloheximide (Sigma-Aldrich 01810) was dissolved in fish water and used at 200 *μm*. During pharmacological treatment, fish were maintained off the aquarium system in the dark. Treatment medium was exchanged every 12 hours.

### Data analysis

Image processing and data analysis were performed with custom-written MATLAB (Mathworks) 2023b software, unless stated otherwise.

#### Preparation of confocal images

For a given regenerating ray, each acquired z-stack contains a portion of the regenerate. Z-stacks tiling each ray were acquired such that each stack contains regions of overlap with the immediate proximal and distal z-stacks. Individual z-stacks were stitched together into a single z-stack spanning the entire proximodistal axis of the regenerate using a custom-written 3D stitching software based on image cross-correlation(De Simone, Evanitsky et al. 2021). These aggregate z-stacks were then rotated so that the entire osteoblast tissue was as parallel as possible to the x-y axes. H2A-mCherry or H2A-mEos2 (non-converted) nuclear signal was used as a nuclear osteoblast reference. As regenerating fin bones mature, osteoblasts surround the ossifying bone of individual hemi-rays. In confocal z-stacks, this is captured as two osteoblast tissue layers. Here, we imaged and analyzed only the external layer of osteoblasts, positioned closest to the overlying epithelium. Fin ray osteoblast tissues were computationally flatted such that the external layer of osteoblast tissue occupied similar z-positions and entire osteoblast tissues where segmented from the nuclear channel images via intensity-based thresholding and morphological operations (De Simone, Evanitsky et al. 2021). The external layer of osteoblasts was computationally isolated (De Simone, Evanitsky et al. 2021). Z-stacks were equalized by contrast-limited adaptive histogram equalization for nuclei segmentation and visualization (De Simone, Evanitsky et al. 2021). Nuclear segmentation was performed on *osterix:H2A-mCherry* or *krt19:H2A-mCherry* signal using TGMM software (Amat, Lemon et al. 2014). Non-equalized z-stacks were used in the quantifications described below.

#### Quantification of photoconversion experiments

Nuclei with photoconverted H2A-mEos2 were manually segmented based on fluorescence intensity in the converted (red) channel. Manual segmentation was confirmed with ratiometric images comparing the converted (red) channel to the unconverted (green) channel. The change in number of nuclei with converted H2A-mEOS was determined by manually counting the number of nuclei with converted fluorescence immediately after photoconversion and 24 hours after photoconversion. The area occupied by photoconverted nuclei was determined by manually drawing the smallest polygon that enclosed all identifiable nuclei with converted fluorescence. To calculate expansion rate, all pixels within a manually defined photoconverted domain were ranked based on their position along the proximodistal axis. The domain edges were defined as the positions of the 10^th^ and 90^th^ percentile of photoconverted pixels. This method ensured that isolated cells did not impact the analysis. Varying the threshold percentiles did not significantly impact the results. Once the edges of the domains were defined, we used their relative distance to compute an average expansion rate for each region: The expansion rate is determined by the divergence of the velocity field, which was estimated as ratio between the difference of the distance between the two edges at the two time points divided by the distance at the initial time point:

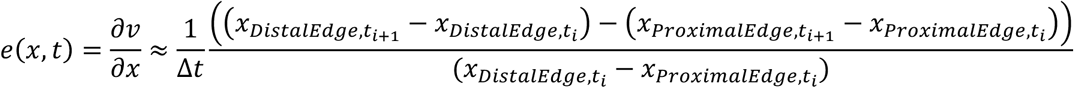

By defining Δ*x* = *x_Distal Edge_* – *x_proximal Edge_*, the previous equation can be recast as:

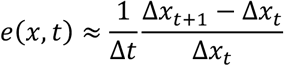

#### Ray length measurements

Length amputated was measured by opening low-magnification, non-compressed TIFF files of caudal fins captured immediately before and after amputation in FIJI. Segmented lines were manually drawn from a fiduciary mark in the post-amputation image to the amputation plane and from the same fiduciary mark in the pre-amputation to image to the distal tip of the rays. The difference between the length of these two lines was recorded as the length amputated. The length of regenerating rays at indicated time points was measured on either stitched confocal images or on low-magnification dissection scope images. For confocal images, regenerate length was defined as the distance between a user-specified distal tip and amputation plane. For dissection scope images, non-compressed TIFF files were opened in FIJI, and segmented lines were drawn along the regenerating rays from the same fiduciary mark used in the post amputation image to the distal tip of the regenerate. Fraction regeneration was calculated as the length of the regenerate at a given experimental timepoint divided by the length of tissue amputated.

#### Identifying and quantifying cycling osteoblasts

To quantify Venus-hGeminin signal in osteoblasts, the nuclear segmentation was used to generate a mask of each nucleus. The nuclear mask was then dilated, and a cytoplasmic mask was generated by subtracting the nuclear mask from the cytoplasmic mask (De Simone, Evanitsky et al. 2021). The amount of *osterix:Venus-hGeminin* expression in each osteoblast was quantified as the average per pixel fluorescence signal in that cell’s nuclear mask normalized by the average per pixel fluorescence signal in that cell’s cytoplasmic mask. Osteoblasts with nuclear:cytoplasmic ratios above a threshold of 1.5 were scored as cycling (unless otherwise noted). Number of osteoblasts cycling (as in Figure 1J) was calculated as the total number of osteoblasts scored as cycling. Fraction osteoblasts cycling (as in Figure 1I, K) was calculated as the total number of osteoblasts scored as cycling divided by the total number of segmented osteoblasts for a given ray. To estimate the length of the proliferative zone (Figure 1L), we plotted the percent Geminin positive of osteoblasts as a function of position along the proximodistal axis and fit these data with a function: *H*(*λ* − *x*)(*b* − *ax*) + *H*(*x* − *λ*)(*b* − *aλ*), where H is the Heaviside step function, *λ* is the size of the proliferative domain, and *x* is the distance from the tip.

#### Erk activity quantification

To quantify Erk activity, the nuclear segmentation was used to generate a mask of each nucleus. The nuclear mask was then dilated, and a cytoplasmic mask was generated by subtracting the nuclear mask from the cytoplasmic mask (as in De Simone et al. 2021). The average per pixel Erk-KTR-mCerulean fluorescence was calculated in the nuclear and cytoplasmic regions, and Erk activity was determined as the ratio of cytoplasmic to nuclear average Erk-KTR-mCerulean signal minus 0.8, which is the average Erk activity value seen in PD03-inhibited fish. To test for oscillatory activity (Supplemental Figure 1C), regenerating rays were imaged every ∼8 hours and divided into 50um long bins, tiling the proximodistal axis. The average Erk activity level within that spatial bin was determined and plotted as a function of time. These spatial bins were aligned at the amputation plane across timepoints for a given ray. We determined the period of Erk activity oscillations by assessing the autocorrelation of Erk activity within spatial bins across time (Supplemental Figure 1D). To determine the distance over which Erk activity is correlated, we performed cross-correlation analysis of Erk activity across varying distances at fixed time (Supplemental Figure 1E).

#### Assessing the relationship between Erk activity and osteoblast cycling

To assess the relationship between Erk activity and osteoblast cycling (as shown in Figure 2G), we imaged regenerating rays every 12 hours for 2-3 days at a time. To overcome the noisy oscillations identified in Supplemental Figure 1C-D and to account for the potential delay between peak Erk activity and Geminin expression, we time averaged Erk activity and fraction cycling values (calculated as described above) across a 24-hour window, which included 3 imaging time points. At each timepoint, rays were divided into 10 equally sized spatial bins tiling the proximodistal axis. Cells within spatial bin 1 at all three timepoints were aggregated, and the average Erk activity and the fraction of osteoblasts cycling was determined. This data point was plotted on Figure 2G, and this approach was repeated for all ten spatial bins. (See also Supplemental Figure 1F).

A modified approach was taken to assess the relationship between Erk activity and osteoblast cycling in *lof* fish (as shown in Figure 5D). Regenerating rays were imaged every 48 hours from 3-21 days post-amputation. At each timepoint, rays were divided into 10 equally sized spatial bins tiling the proximodistal axis, and the average Erk activity within a given spatial bin was plotted against the fraction of osteoblasts cycling in that spatial bin at a single timepoint. The red line in Figure 5D is the average of two independent experiments in *lof* fish.

#### Whole ray average Erk activity

Whole ray average Erk was quantified by averaging the Erk activity of all cells within a regenerating ray at a single time point.

#### Spatial profiles of Erk activity

Spatial profiles of Erk activity, such as those seen in Figure 4B and Supplemental Figure 4B, were generated by dividing rays into spatial bins of 250 *μm* calculating the average Erk activity of osteoblasts within each spatial bin and plotting that as a function of absolute position along the proximodistal axis of regenerates. Normalized spatial profiles, such as those seen in Supplemental Figure 4C, were generated by dividing rays into 10 equally sized spatial bins, calculating the average Erk activity within each spatial bin normalized by the whole ray average Erk, and plotting these values against the normalized position along the proximodistal axis. To generate the representative Erk activity curves shown in Figures 4C, Figure 4D, Figure 5E, and Supplemental Figure 5D, we separated the normalized spatial profiles into five groups according to each ray’s length amputated (Figure 4C, 5E) or time post amputation (Figure 4D, Supplemental Figure 5D). We then calculated the average normalized Erk activity profile of all rays within a given length amputated or time post amputation group.

To generate the representative Erk activity profile throughout regeneration (shown in Figure 4E), we first generated Erk activity profiles as a function of normalized regenerate position for each ray by creating 10 equally sized bins along the regenerate, spanning from distal tip to amputation plane. The average Erk activity within each spatial bin was determined as a function of normalized regeneration position (x / length regenerate). These Erk activity versus normalized regenerate position profiles were then grouped into 10 groups according to their fraction regeneration (i.e. the length regenerated / length amputated at the time point analyzed). For example, if a ray had a fraction regenerated of 0.25, it would be grouped into the 0.2 <= phi < 0.3 group. The mean of all Erk activity versus normalized regenerate position profiles was then calculated for all 10 groups. Finally, this mean Erk versus normalized regenerate position was plotted along the x axis starting at 0 (representing the amputation plane) and ending at the median phi value of a given group.

To generate a quantitative description of Erk activity in space *f(u)*, the normalized Erk activity versus normalized position profiles of all analyzed rays were collected and fit with line.

#### Quantifying tissue growth, Erk activity, and osteoblast cycling during pharmacological treatments

Confocal images of rays expressing Erk KTR-mCerulean, Venus-hGeminin, and/or H2A-mCherry were acquired and stitched, as described above, before and after pharmological treatment. Tissue growth (as in Figure 2B & 6D) was calculated by subtracting regenerate length before pharmacological inhibition from the regenerate length after pharmacological inhibition. The effect of pan-Fgf Receptor inhibition on Erk activity was assayed by comparing the whole ray average Erk activity level in individual rays before and after pan-FgfR inhibition (Figure 6B). The effect of pharmacological treatment on osteoblast cycling was assayed by determining the number of osteoblasts scored as cycling before and after pharmacological inhibition (Figure 2D & 6C)

#### Quantification of fgf20a:EGFP expression

To quantify GFP expression from Fgf20a regulatory elements, images of fin rays expressing *fgf20a:*EGFP and *krt19*:H2A-mCherry were cropped such that they contained only one bony ray, extending from the amputation site to the distal tip of the epidermis. Epidermal tissue immediately surrounding the bone dorsally and ventrally was included. Epidermal nuclei in the cropped images were segmented on signal from krt19:H2A-mCherry using TGMM. Since *fgf20a:*EGFP signal is distributed uniformly between the nucleus and cytoplasm, the average per pixel *fgf20a:*EGFP expression level per cell was estimated as the average GFP intensity per pixel within the nuclear mask generated during segmentation. Cells were scored as *fgf20a:*EGFP expressors if the average nuclear fluorescence signal was greater than 50 A.U.

To quantify the length of the *fgf20a*:EGFP expression domain, the GFP signal of each cropped z-stack was integrated along the Z and Y axes of the image. The proximal edge of the *fgf20a:*EGFP expression domain was then determined as the X position at which GFP intensity fell below 10.5% of the max GFP intensity value for a given ray. The length of the *fgf20a:*EGFP expression domain was determined as the distance between the distal tip and the proximal edge.

For the experiments quantified in Figure 7G (blue data) and Supplemental Figure 7B,C, we were unable to measure the regenerate length at the regeneration end point (∼3-4 weeks). Instead, we estimated the final length of each WT or lof ray using the growth dynamics observed for WT or *lof* fish (Supplemental Figure 5B&C, respectively).

### Theoretical model for the formation and evolution of Erk gradients

#### Limit of negligible diffusion

In the main text, we have introduced a mathematical model describing a signaling gradient forming by displacement of ligands due to molecular diffusion and tissue growth. The model takes the following form:

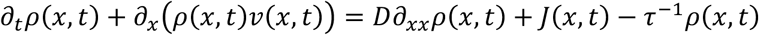

Various arguments suggest that diffusion is negligible in this system (see main text and below).

In the limit of negligible diffusion, the model equation reduces to:

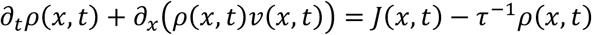

This equation can be solved mathematically using the method of the characteristics, which is equivalent to computing how the concentration evolves along lagrangian trajectories:

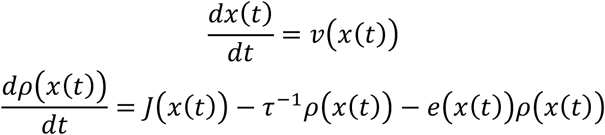

where *e*(*x*(*t*)) = 𝜕*_x_v*(*x*(*t*)) is the rate of expansion of the tissue that determines the dilution of the ligand. Intuitively this framework can be thought of as a mathematical way to describe the movement of individual cells, captured by the lagrangian trajectories obtained solving the first equation, and compute how the concentration of ligands seen by these cells changes over time. To determine if this model can reproduce the experimentally observed Erk gradients, one can compute *ρ*(*x*, *t*) for a given source *J*(*x*, *t*) and lifetime of the ligand *τ*. However, since these parameters are difficult to evaluate quantitatively, we decided to invert the relationship between the dynamics of the ligand and the dynamics of the source to obtain:

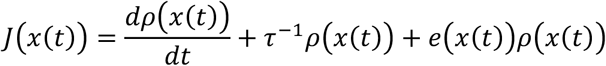

We then compute the spatiotemporal dynamics of the source that reproduce the observed gradients for a given value of the ligand lifetime. To do this, we assumed that the concentration of ligand is proportional to Erk activity and took 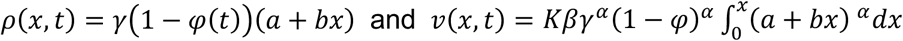, see main text for details. Thus, we can directly compute all the terms on the right hand-side for each position in the ray once a value of *τ* is specified. We also used the expression:

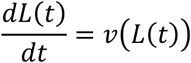

to compute the dynamics of tissue elongation. The differential equations were solved numerically using the ode45 solver in MATLAB. This analysis shows that a transient and localized source of ligands, qualitatively similar to the spatiotemporal properties of *fgf20a:*EGFP expressing cells, and a ligand lifetime *τ* of about 4 days or longer can generate the dynamic, scaling gradients of Erk activity observed experimentally (see Figure 6). The long lifetime of the ligand is supported by experiments inhibiting protein translation with cycloheximide support this ligand lifetime value:

Erk activity decays at a similar rate in control and treated animals over a 12-hour period (longer treatment were toxic), suggesting that ligand lifetime is likely to be significantly longer than a day (Figure 6H). Furthermore, measurements of the temporal decay of Erk activity near the amputation plane, where we assume that there is little to no contribution from ligand production or movement, yield an estimated lifetime of 3.7 days, a value ultimately used in our simulations and close to our computationally estimated value (Supplemental Figure 7E).

Having established that the previous model can explain the formation of dynamic gradients from a transient and localized source, we turned our attention to understanding the mechanisms controlling scaling and positional memory (control of regenerate tissue size) in this model. We note that the only length scale present in the model is the size of the source. Thus, we reasoned that if the properties of the source scales with the amount of tissue amputated that should be sufficient to ensure positional memory, which was confirmed by simulations (Supplemental Figure 6). Thus, our theoretical analysis leads to a simple model for the encoding of positional memory by the number of ligand-producing cells in the source.

#### Arguments against a major role for diffusion

Gradients could in principle form by diffusion and degradation, as observed and/or proposed for other morphogen gradients (Bollenbach, Kruse et al. 2007, Grimm, Coppey et al. 2010, Kicheva, Cohen et al. 2012). This is equivalent to ignoring the advection/dilution term, 𝜕*_x_*(*ρ*(*x*, *t*)*v*(*x*, *t*)) in the original equation. For a diffusion constant *D* and a lifetime of the ligand *τ*, the length scale of the gradient at steady state is given by: 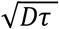. *τ* also sets the time needed for the gradient to reach steady state. We argue that it is unlikely that a combination of diffusion and degradation is what drives the formation of the Erk signaling gradients in this system (see also main text). This is based on three main observations. First, the previously inferred value of the diffusion constant of Fgf ligand(s) implicated in osteoblasts regeneration (*D* ≈ 0.1*μm*^2^/*s*) would imply a value of *τ* ≈ 1 *year* for gradients of about 2 millimeters in length as those observed in mid-regeneration in lateral rays. Such a long lifetime of the ligand is incompatible with a regeneration process lasting about 3 weeks. Second, diffusion is expected to be isotropic. If diffusion was able to spread ligands across millimeters not only proximo-distally but also laterally, then the dynamics of most rays would be coupled (the typical inter-ray distance is about 400 *μm*), which is inconsistent with evidence that the regeneration dynamic of each ray is independent from others (Owlarn, Klenner et al. 2017, Uemoto, Abe et al. 2020). Third, the experimentally observed scaling of gradients: *λ* ≈ *L* and the large range over which scaling is observed (gradient length scales ranging between 0.2 − 4 *mm*) would implicate that the diffusion constant and/or lifetime must be precisely fine-tuned across two orders of magnitude in both space and time. No such fine-tuning of parameters is required for the advection model we presented.

### Bulk RNAseq

To isolate RNA from the amputation plane, a region spanning one bone segment above and below the amputation plane was collected from fish at 0, 3.5, 7, 14, and 21 dpa. For each time point, tissue samples from 10 animals were pooled for each of 3 biological replicates and collected in Tri-Reagent (Sigma). To isolate RNA from the distal tip of regenerating rays, a region spanning the distal ∼250 µm of each ray was collected from fish at 0, 3.5, 7, 10, and 14 dpa. For each time point, tissue samples from 10 animals were pooled for each of 3 biological replicates and collected in Tri-Reagent (Sigma). Only 2 biological replicates were of high enough quality for bulk RNAseq. Tissue was homogenized with a stainless-steel bead (5mm, Cat. No. 69989) in Tri-Reagent. RNA was extracted by cholorform followed by an isopropanol precipitation. Remaining DNA was digested by DNase. RNA was purified with a Zymo RNA Clean & Concentrator Kit. For tip tissue samples, library preparation was performed by BGI Genomics. For amputation plane tissue samples, cDNA was prepared by Maxima H-minus RT reverse transcription kit. PCR was performed using Kapa HiFi HotStart Master Mix and adapter primers. The amplified cDNA was purified using 0.6x SPRI beads. cDNAs were tagged using Tn5 transposase and amplified by Q5 polymerase. Size selection was performed with gel-cutting and extraction, and 300 - 800 bp DNA fragments were collected for sequencing. Libraries were sequenced using a DNBSEQ-MGI2000, 150 bp PE, by BGI Americas. RNA-seq reads were trimmed by Trim Galore (v 0.6.7, with cutadapt v 3.5) and counted by salmon (v 1.4.0) by supplying the UCSC danRer11 annotation (Patro, Duggal et al. 2017). Bioconductor package DESeq2 (v 1.34.0) (Love, Huber et al. 2014) was employed to analyze differential expressions (DE) with genotype information.

---

**Supplemental Figure 1.**
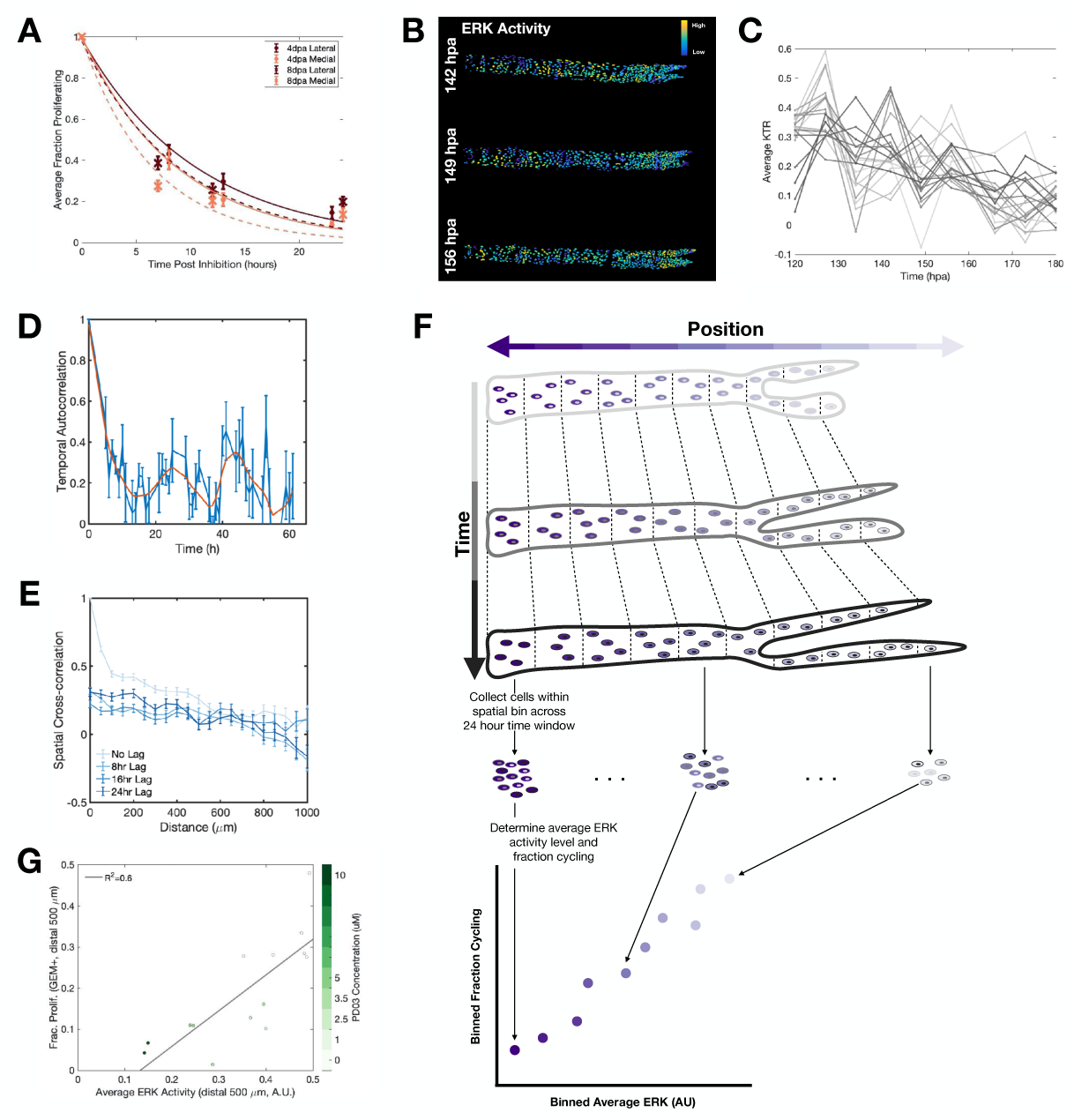
Erk activity oscillates and quantitatively influences cell cycling in osteoblasts. A) Quantification of the fraction of osteoblasts expressing Venus-Geminin as a function of time over the course of a 24 hour MEK inhibitor (PD03) treatment. Erk activity is required for the G1 to S transition, so no new cells are expected to enter the cell cycle during PD03 treatment. Thus, the osteoblast cell cycle during fin regeneration can be estimated as the time it takes for Venus-Geminin expression to cease. Here, we estimated the osteoblast cell cycle to be ∼24 hours. For each curve, 4 rays from 4 fish were scored. (A total of 16 rays from 16 fish were scored.) B) Representative heat map of regenerating ray expressing *osterix*:Erk-KTR-Cerulean and *osterix*:Histone-2A (not shown) imaged ∼every 8 hours. Osteoblast nuclei are color coded according to their Erk activity level. C) Quantification of Erk activity along the proximodistal axis of the regenerating ray shown in (B) as a function of hours post amputation. Each line represents a distinct 50 *μm* wide spatial bin tracked throughout time. Data is plotted in grayscale to distinguish individual traces. 19 bins from 1 ray are plotted. D) Autocorrelation analysis to test for temporal oscillations. Noisy temporal oscillations with a period of ∼24 hours were identified. Data are from 10 rays from 5 fish. E) Cross-correlation analysis to test for traveling or standing waves. No evidence of traveling or standing waves was observed. Data are from 10 rays from 5 fish. F) Explanation of Quantification of Binned Fraction Cycling versus Binned Erk Activity: Time is shown along the vertical axis and is depicted by gray shading of nuclei. Proximodistal position is shown along the horizontal axis and is depicted by purple shading of cytoplasm. We imaged regenerating rays every 12 hours for 2-3 days at a time. To overcome the noisy oscillations identified in Supplemental Figure 1C and to account for the potential delay between peak Erk activity and Geminin expression, we time averaged Erk activity and fraction cycling values across a 24-hour window, which included 3 imaging time points. At each timepoint, rays were divided into 10 equally sized spatial bins tiling the proximodistal axis. Cells within spatial bin 1 at all three timepoints were aggregated, and the average Erk activity and the fraction of osteoblasts cycling was determined. This was repeated for all ten spatial bins, and this data point was plotted on Figure 2G. G) Quantification of fraction osteoblasts cycling versus average binned Erk activity at the distal tip rays of MEK inhibitor (PD03) treated fish. Each dot represents a spatial bin spanning the distal-most 500um of a regenerating ray, colored according to the concentration of MEK inhibitor (PD03) with which the fish was treated. 4 fish/rays were treated with DMSO control, 2 fish/rays each were treated @ 1 *μm*, 2.5 *μm*, 3.5 *μm*, 5 *μm*, and 10 *μm*. (A total of 14 fish/rays were used.)

**Supplemental Figure 2.**
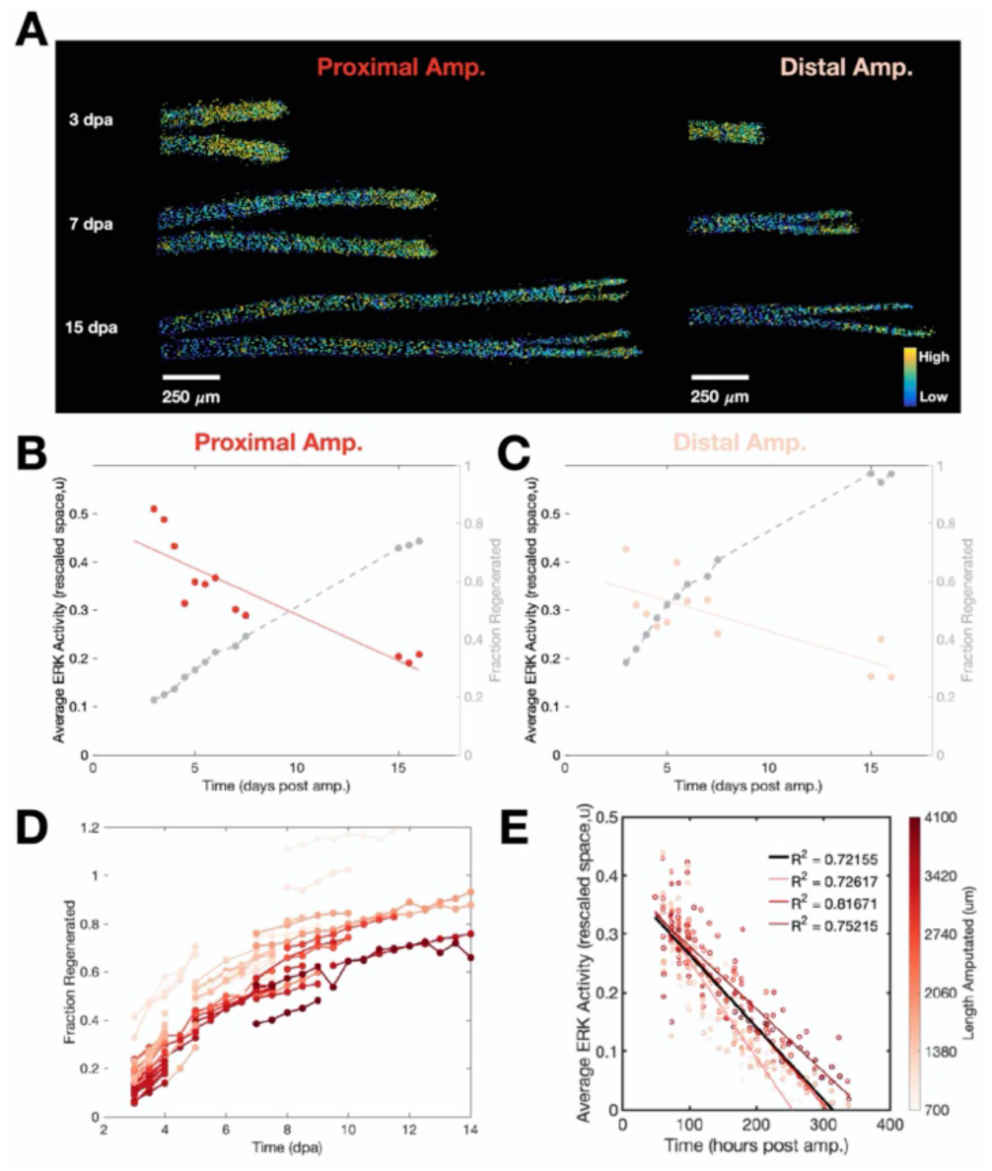
Whole-regenerate averaged Erk activity decreases over time in individual regenerating rays. A) Representative heatmaps of Erk activity in regenerating rays from proximal (left) and distal (right) amputations. (Reproduced from Figure 3A in main text.) B,C) Quantification of whole ray average Erk activity (red) and fraction regenerated (gray) versus time post amputation for representative rays shown in Supplemental Figure 2A. Red lines are fits to average Erk activity versus time data. D) Quantification of fraction regenerated over time for all analyzed rays. Lines connect data points from the same ray. Coloring indicates length amputated. Data are from 60 rays from 22 fish. E) Whole ray average Erk activity versus time. Each dot represents a single ray at a single time point, colored according to length amputated. Red lines indicate fits data sub-grouped by amputation length. Black line indicates fit to all data. Data are from 60 rays from 22 fish.

**Supplemental Figure 3.**
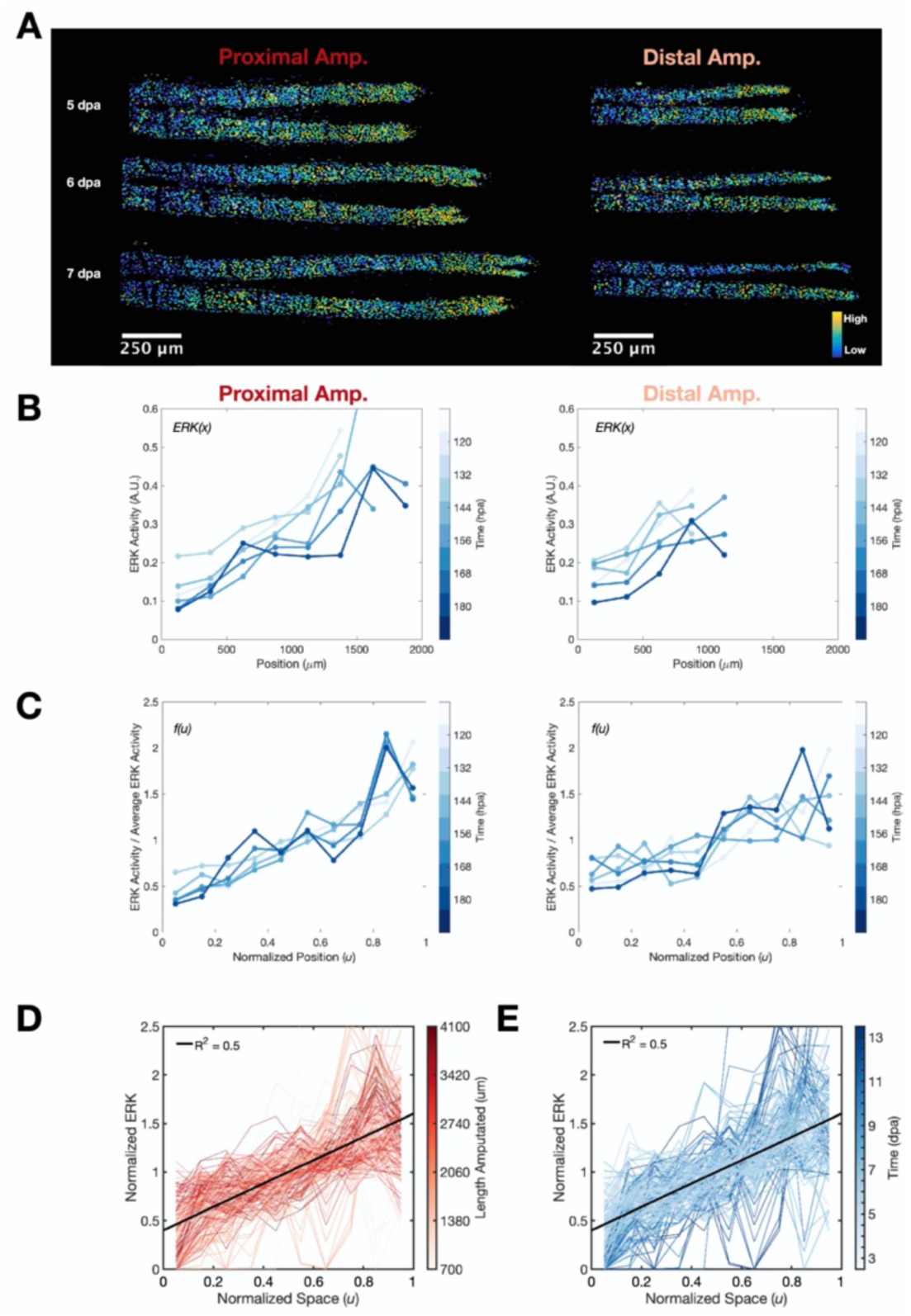
Normalization of spatial Erk activity patterns in space and time. A) Representative images of Erk activity heatmaps for regenerating rays with proximal (left) or distal (right) amputations, imaged every 12 hours from 5 to 7 days post amputation. (Repeated from Figure 4A in main text.) B) Quantification of Erk activity along the proximodistal axis the regenerating rays shown in Figure 4A. Amputation site is plotted at 0 on x-axis. Each dot represents the average Erk activity of cells within a spatial bin. Lines connect points from the same ray at a given time. Coloring indicates time post amputation. (Repeated from Figure 4B in main text.) C) Data as in Supplemental Figure 3B except y-axis values are normalized by whole ray Average Erk and x-values are normalized by length regenerated to facilitate comparisons between rays at different timepoints. Coloring indicates time post amputation. D) Plot of normalized Erk activity (Binned Average Erk activity / Whole ray average Erk activity, see Supplemental Figure 3C for example of single ray traces) versus normalized position along the proximodistal axis. Each line represents a single ray whose spatial Erk activity measurements were averaged over a 24 time window (See Supplemental Figure 1F for more details). Coloring indicates length amputated. Black line indicates fit to all data. Data are from 52 rays from 20 fish. E) Plot as in Supplemental Figure 3D with data colored by time post amputation instead of length amputated. Data are from 52 rays from 20 fish.

**Supplemental Figure 4.**
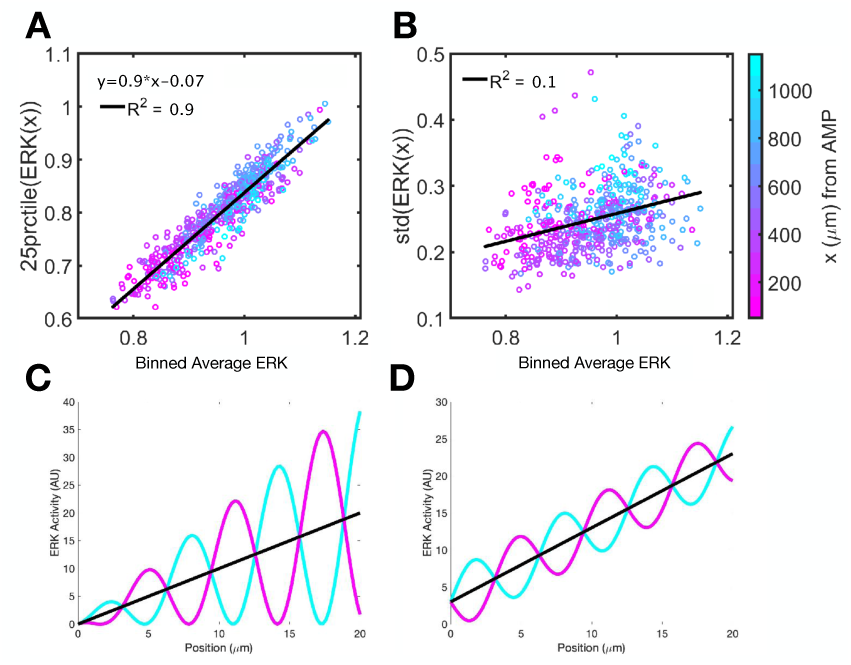
Oscillations of Erk activity of similar amplitude across space overlie on long-range gradients. A) Plot of mean Erk activity of the 25^th^ percentile of Erk values for a given spatial bin versus the mean of all Erk values in that bin. Each dot represents a spatial bin, colored by distance from amputation plane. The observation that the minimum Erk activity increases as average Erk activity increases supports the outcome diagrammed in Supplemental Figure 4D B) Plot of the variance of the Erk values within a given spatial bin versus the mean of the Erk values in that bin. Each dot represents a spatial bin, colored by distance from amputation plane. The observation that the variance in Erk activity remains constant along the length of the regenerate supports the outcome diagrammed in Supplemental Figure 4D. C) Schematic of oscillations whose minima are similar along the proximodistal axis but whose amplitudes increase toward the distal tip. Colors indicate two distinct time points. D) Schematic of oscillations whose amplitudes are similar but whose averages gradually increase toward the distal tip. Colors indicate two distinct time points.

**Supplemental Figure 5.**
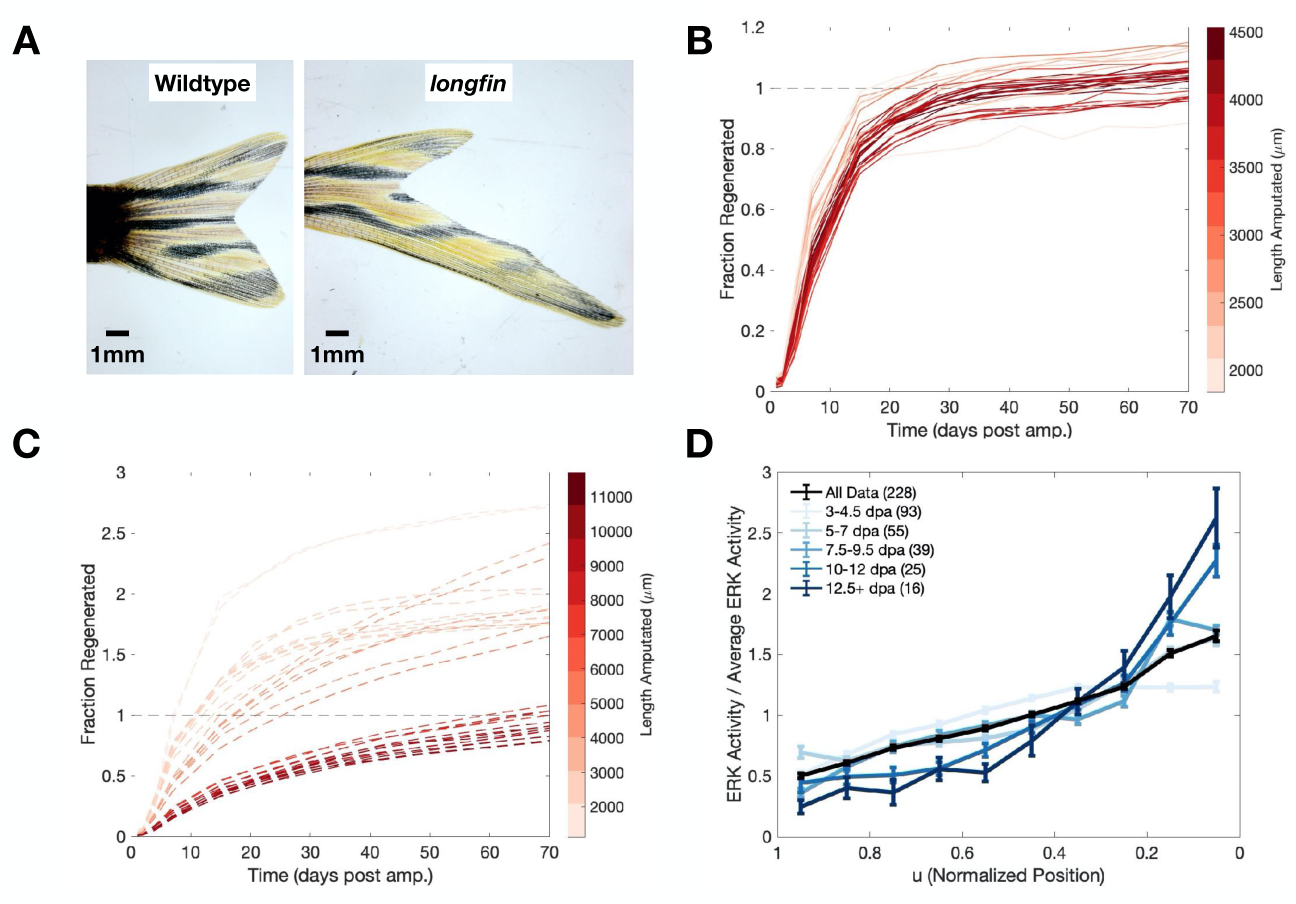
Size memory is compromised in *longfin* mutant fish, but *Erk* activity persists as gradients throughout fin regeneration. A) Representative images of uninjured fins from wildtype (left) and *lof* mutant (right) fish. B) Quantification of regenerative growth dynamics in wildtype fish. Data are from 36 rays from 6 fish. C) Quantification of regenerative growth dynamics in *lof* fish. Data are from 30 rays from 5 fish. D) Spatial Erk activity profiles of *lof* rays, as in Figure 5E, colored by time post amputation instead of length amputated. Data are from 36 rays from 9 fish.

**Supplemental Figure 6.**
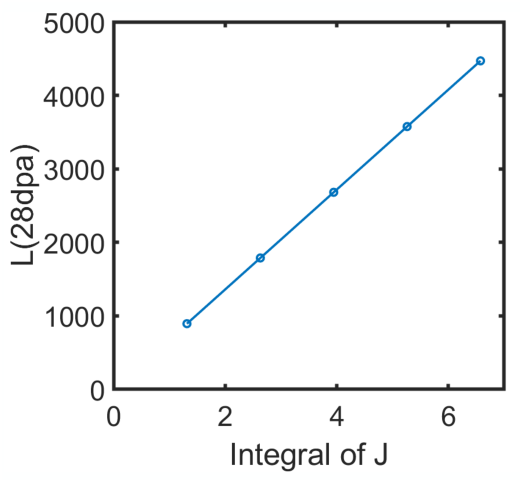
The length regenerated is explained by the extension of the source domain. Comparison of the extension of the source domain (J) to the length predicted by our model to be regenerated at 28 days post amputation.

**Supplemental Figure 7.**
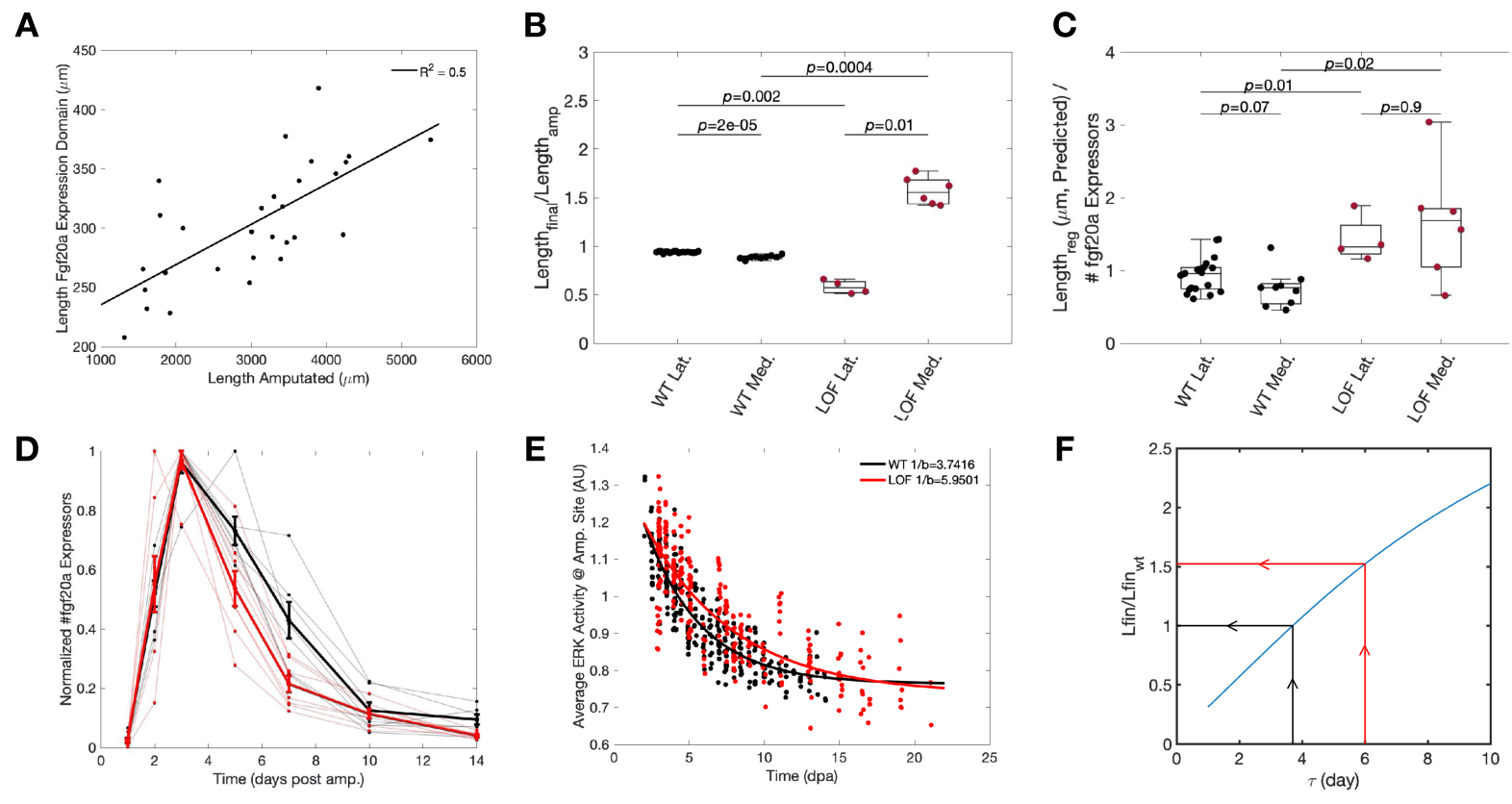
Erk activity decays more slowly in *longfin* mutant fish. A) Quantification of the length of the *fgf20a:*EGFP expressing domain at 72 hpa versus length amputated. Each dot is a single regenerating ray. Black line is fit of all data. Data are from 41 rays from 16 fish. B) Quantification of the ratio between length amputated and final predicted length for rays analyzed in Figure 7. Black – wildtype data, red – *lof* data. Wildtype data are from 26 rays from 12 fish. *lof* data are from 10 rays from 4 fish. C) Quantification of ratio between final predicted length and number of *fgf20a:*EGFP expressors for rays analyzed in figure 7. Black – wildtype data, red – *lof* data. Wildtype data are from 26 rays from 12 fish. lof data are from 10 rays from 4 fish. D) Quantification of *fgf20a:*EGFP expression dynamics, as scored by *fgf20a*:EGFP expression, in wildtype (black) and *lof* (red) fish. Bold lines indicated average dynamics for each genotype. Wildtype data are from 7 rays from 4 fish. lof data are from 8 rays from 4 fish. E) Quantification of Erk activity decay in a region spanning 0-250 *μm* from the amputation site in wildtype (black) and *lof* (red) fish. Estimated ligand lifetimes are shown in inset. Data are from 60 rays from 22 fish. F) Ratio of length regenerated in *lof* to length regenerated in wildtype fins predicted by mathematical model for different ligand lifetime values in *lof* fish. A ligand lifetime of ∼6 days yields 1.5 times more growth than a wildtype ray (red lines), consistent with the ligand lifetime estimated for *lof* fish in Supplemental Figure 7E.

## Acknowledgements

We thank J. Burris, K. Oliveri, C. Dolan, L. Frauen, D. Stutts, K. Scallion, M. Powell, and M. Wiley for zebrafish care. We thank B. Hogan and B. Mathey-Prevot for scientific discussions, advice, and critical reading of the manuscript. This work was supported by N.I.H grants 5F32HD107853 to A.R., R01 AR076342 to K.D.P. and S.D., and R01 HD105033 to K.D.P. It was also supported by an Innovation in Stem Cell Science Award from the Shipley Foundation, Inc. to S.D.

